# AlphaFold3-based modeling uncovers the dynamic structural interface between full-length IAP antagonists and DIAP1 for apoptosis regulation in *Drosophila*

**DOI:** 10.64898/2026.01.09.698746

**Authors:** Pooja Rai, Andreas Bergmann

## Abstract

Apoptosis in *Drosophila* is governed by caspases, inhibitor of apoptosis proteins (IAPs), and IAP antagonists. Using AlphaFold3, we modeled full-length 3D structures of the IAP antagonists Reaper, Hid, Grim, Sickle, and Jafrac2, as well as DIAP1 and dBruce, and their binary and higher-order complexes. We uncover a paradoxical role for the N-terminal methionine of Reaper in stabilizing Reaper/Hid complexes and inhibiting DIAP1 binding. Our models reveal that Reaper uniquely engages both BIR1 and BIR2 domains of DIAP1, guided by α-helical residues in its backbone, while all other IAP antagonists preferentially target BIR2. Higher-order assemblies show how Reaper and Hid cooperatively engage DIAP1 and allosterically modulate its E3 ligase activity. We present the first full-length model of dBruce and its inhibitory interaction with Rpr. These findings provide a comprehensive structural framework for apoptosis regulation in *Drosophila*, and offer new insights into conserved mechanisms of caspase control and IAP antagonism across species.

## 1. Introduction

Apoptosis is a genetically programmed form of cell death characterized by distinct morphological and biochemical features, including cell shrinkage, chromatin condensation, and DNA fragmentation (Kerr et al., 1972). It serves as a fundamental mechanism to eliminate superfluous, damaged, or potentially dangerous cells during development and tissue homeostasis. Dysregulation of apoptosis can lead to a wide array of human diseases, including cancer, neurodegeneration, and autoimmune disorders (Thompson, 1995).

A hallmark of apoptosis is the activation of caspases, a family of cysteine proteases that cleave cellular substrates at aspartate residues to execute the death program (Green, 2024; McIlwain et al., 2013; Yuan and Ofengeim, 2024). While caspase activation lies at the heart of apoptosis, its regulation is tightly controlled by Inhibitor of Apoptosis Proteins (IAPs), which directly bind and inhibit caspases to prevent inappropriate cell death (Gyrd-Hansen and Meier, 2010; Vaux and Silke, 2005). IAPs are evolutionarily conserved and defined by the presence of one or more Baculovirus IAP Repeat (BIR) domains. Several IAPs also contain a C-terminal RING domain that confers E3 ubiquitin ligase activity. While mammals encode at least seven IAPs, *Drosophila* possesses four. In *Drosophila*, the central IAP is DIAP1 (*Drosophila* IAP1), which functions both as a caspase inhibitor and as an E3 ubiquitin ligase that targets caspases and itself for ubiquitylation. Loss of DIAP1 causes extensive apoptosis, demonstrating its essential role in cell survival (Goyal et al., 2000; Lisi et al., 2000; Wang et al., 1999).

Apoptosis in *Drosophila* is further controlled by a family of pro-apoptotic proteins known as IAP antagonists. The best-characterized are Reaper (Rpr), Head involution defective (Hid), and Grim, which are clustered in the genome and whose combined deletion results in complete elimination of apoptosis (Chen et al., 1996; Grether et al., 1995; White et al., 1994). A fourth member, Sickle (Skl), was identified based on sequence similarity to Rpr (Srinivasula et al., 2002; Wing et al., 2002), though its physiological role remains less well defined. A fifth IAP antagonist, Jafrac2, is a thioredoxin peroxidase encoded by the *Prx4* gene (Tenev et al., 2002). Unlike other IAP antagonists, Jafrac2 is localized to the endoplasmic reticulum (ER) and functions in oxidative protein folding under basal conditions. Upon apoptotic signaling, Jafrac2 is released into the cytosol where it antagonizes DIAP1 (Tenev *et al*., 2002).

The defining feature of IAP antagonists is the presence of a short N-terminal IAP-binding motif (IBM) (typically Ala-Val/Ile-Ala/Pro-Phe-Tyr) (Chen *et al*., 1996; Grether *et al*., 1995; Vucic et al., 1998; Wing et al., 2001), which is exposed after removal of the initiator methionine (Wu et al., 2000). This motif enables direct binding to the BIR domains of IAPs, displacing caspases and promoting cell death. Rpr and Grim also contain a central α-helical Grim Helix 3 (GH3) domain (Claveria et al., 2002; Wing *et al*., 2001), which promotes their multimerization and mitochondrial localization (Olson et al., 2003; Sandu et al., 2010), two features which are essential for full apoptotic activity. Hid lacks the GH3 motif but contains a transmembrane tail-anchor (TA) domain at its C-terminus that targets it to the outer mitochondrial membrane (OMM). The TA domain of Hid also serves as a platform to recruit other IAP antagonists such as Rpr to the mitochondria, where higher-order complex formation is essential for apoptosis in *Drosophila* (Sandu *et al*., 2010). Rpr, in particular, requires Hid for efficient mitochondrial localization and apoptotic activity. Both mitochondrial targeting and forced dimerization of Rpr strongly enhance its pro-apoptotic function (Sandu *et al*., 2010).

dBruce is another IAP family member that plays a distinct role in apoptosis regulation (Arama et al., 2003; Vernooy et al., 2002). It contains an N-terminal BIR domain and a C-terminal Ubc domain with E2-like ubiquitin conjugating activity. Genetically, dBruce has been shown to inhibit Rpr-induced apoptosis, possibly by sequestering Rpr and promoting its degradation through an unconventional ubiquitination mechanism (Domingues and Ryoo, 2012; Vernooy *et al*., 2002).

To date, most structural insights into IAP-antagonist interactions have come from X-ray crystallography of short IBM peptides (10mers) bound to isolated BIR domains (Chai et al., 2003; Wu et al., 2001; Yan et al., 2004). These X-ray structures provided high-resolution views of how individual motifs engage specific binding pockets in BIR1 or BIR2. However, due to the inherent challenges of crystallizing large, flexible, and often disordered full-length proteins, structural information on full-length IAP antagonists and DIAP1 in their native multimeric states remains limited. Furthermore, isolated domains and short peptides cannot fully capture the cooperative interactions and conformational rearrangements that may occur when entire proteins engage in complex regulatory assemblies.

To address this gap, we employed AlphaFold 3 (AF3), a state-of-the-art deep learning model that predicts protein structures and protein-protein interactions (PPIs) with remarkable accuracy (Abramson et al., 2024; Jumper et al., 2021). AF3 integrates sequence, structural, and evolutionary information to generate accurate models of individual proteins and their complexes, even in the absence of experimental data (Jumper *et al*., 2021). We validated our models using the predicted Local Distance Difference Test (pLDDT) score (Mariani et al., 2013), which provides a confidence estimate for each residue’s position in the predicted structure, and the Predicted Aligned Error (PAE) map which indicates the position error of two residues relative to each other (Abramson *et al*., 2024; Elfmann and Stulke, 2023; Varadi et al., 2022).

Here, we apply AF3 to systematically model the structures of full-length Rpr, Grim, Hid, Sickle, Jafrac2, DIAP1, and dBruce, as well as their binary and higher-order complexes. Our analysis reveals new insights into the structural basis of IAP antagonist function including the unstructured, but functionally dynamic architecture of Hid, the role of dimerization, domain-specific binding, and cooperative inhibition of DIAP1. We identify key residues and motifs that mediate these interactions and uncover how structural context influences binding specificity.

Notably, we show that Rpr binds both BIR1 and BIR2 of DIAP1, while all other IAP antagonists primarily target BIR2. We also explore how higher-order complexes involving Rpr and Hid modulate the E3 ligase activity of the RING of DIAP1, and how dBruce may inhibit Rpr through sequestration and unconventional ubiquitination.

These structural insights provide a more complete picture of the apoptotic machinery in *Drosophila*. Our findings not only validate known biochemical interactions, but also uncover previously unappreciated modes of complex assembly and domain coordination. This study provides a new molecular framework for understanding how *Drosophila* cells regulate apoptosis and offers a rich resource for future structural and functional studies in both invertebrate and vertebrate systems.

## 2. Methodology

### 2.1. Structural Modeling Using AlphaFold 3

To predict the full-length structures of DIAP1, IAP antagonists, and associated apoptotic regulators in *Drosophila melanogaster*, we employed AlphaFold3 (AF3), a state-of-the-art deep learning framework for protein structure prediction (Abramson *et al*., 2024). Protein sequences were obtained from FlyBase, UniProt, and the RCSB Protein Data Bank. AF3 integrates multiple sequence alignments, homologous structural templates, and deep neural networks trained on physical constraints and co-evolutionary signals to generate atomic-resolution models (Abramson *et al*., 2024; Jumper *et al*., 2021). This allowed us to capture not only domain architectures but also inter-domain arrangements and flexible structural elements in both monomeric and multimeric contexts.

### 2.2. Multimer Mode and Model Validation

We utilized AF3’s prediction mode to generate heteromeric complexes comprising IAP antagonists and DIAP1. Unlike the available *Drosophila* predictome, which is based on AlphaFold2 and only predicts individual protein structures, or AlphaFold2 Multimer, which can model protein-protein complexes, but cannot incorporate cofactors (such as Zn^2+^), AF3 allows full multimer modeling with cofactors included. This multimer modeling eliminates the need for sequential docking, offering a powerful and integrative approach to predict native-like macromolecular assemblies.

To assess model reliability, we evaluated per-residue confidence scores (pLDDT) and Predicted Aligned Error (PAE) maps, which estimate positional accuracy within and across subunits (Abramson *et al*., 2024; Elfmann and Stulke, 2023; Jumper *et al*., 2021; Mariani *et al*., 2013). As an internal benchmark, we compared AF3-predicted structures of BIR1 and BIR2 domains of DIAP1 bound to IBM peptides with their available crystallographic complexes (Wu *et al*., 2001). The close agreement in structural features and interface residues validated the predictive accuracy of AF3 and supported its use for modeling full-length complexes which are otherwise not easily amenable to crystallization.

### 2.3. Protein-Protein Docking

Monomeric structures of Rpr, Hid, Grim, Jafrac2, Sickle, DIAP1 and dBruce were first predicted using AF3. To independently evaluate multimeric interfaces and explore binding geometries, we employed two complementary approaches based on the protein complexes under investigation. Docking of heterodimeric IAP antagonist complexes was performed using the ClusPro web server (https://cluspro.bu.edu/). ClusPro performs an FFT-based global search of rotational and translational space to efficiently evaluate millions of possible docking orientations. Energetically favorable poses are subsequently clustered, and top-ranking clusters were visually and energetically analyzed to identify plausible protein-protein interfaces.

Separately, AF3 was employed to model multimeric assemblies including the Rpr homodimer and all protein complexes containing IAP antagonists and DIAP1 with chelated Zn^2+^. All predicted structures were exported in PDB format, and molecular docking was performed using default scoring conditions that integrate electrostatics, desolvation, and van der Waals contributions. Among the resulting models, the cluster with the highest overall confidence score was selected for detailed structureal analysis.

### 2.4. Structural Annotation Using PDBsum

For comprehensive annotation of structural features, predicted complexes were submitted to PDBsum (https://www.ebi.ac.uk/thornton-srv/databases/pdbsum/). PDBsum provides detailed diagrams of secondary structure content, topology, domain boundaries, and interaction schematics, including LigPlot+ interaction maps of non-covalent contacts (Laskowski, 2022; Laskowski et al., 2018). It identifies residues at the interface that are involved in hydrogen bonds, salt bridges and non-bonded contacts. These annotations facilitated analysis of how specific motifs and residues contribute to DIAP1/IAP antagonist binding.

### 2.5. Protein-Protein Interaction Analysis

Detailed analysis of protein-protein interfaces was conducted for DIAP1 in complex with Rpr, Hid, Grim, Sickle, and Jafrac2. AF3-derived complexes were interrogated to identify interacting residues, binding hotspots, and structural determinants of specificity. Emphasis was placed on interactions involving the BIR1 and BIR2 domains of DIAP1, the IAP-binding motifs (IBMs) of IAP antagonists, and additional motifs such as the GH3 helix. This analysis revealed both canonical and novel contacts not captured in peptide-based crystal structures, offering insights into full-length protein binding behavior.

### 2.6. Molecular Visualization

All molecular graphics and visual analyses were performed in PyMOL 3 (Schrödinger, LLC). Structural models were inspected for domain architecture, conformational integrity, and intermolecular contacts. We specifically examined Zn^2+^ coordination sites in BIR and RING domains of DIAP1, IBM docking pockets, the relative orientation of IAP antagonists in complex with DIAP1, and the domain architecture of DIAP1 after binding of IAP antagonists.

Representative figures were generated to highlight key structural features and binding modes, including domain-domain interactions, multimeric assembly topologies, and conformational adaptations. These high-resolution renderings were used for figure panels and to guide biological interpretations of structure-function relationships in the apoptotic regulatory network.

## 3. Results

### 3.1 Structures of the full-length proteins of Reaper, Hid, Grim, Sickle, and Jafrac2 predicted by AlphaFold3

To gain structural insights into the regulation of apoptosis in *Drosophila*, we used AlphaFold 3 (AF3) to predict the full-length structures of the IAP antagonists Rpr, Hid, Grim, Sickle, and Jafrac2. These proteins promote cell death by antagonizing DIAP1, primarily through their N-terminal IAP-binding motifs (IBMs).

Rpr encodes a small protein of 65 amino acids (aa) (White *et al*., 1994). The predicted structure reveals a central elongated α-helix spanning residues 10-50, which includes the GH3 motif (aa 32-46) (Figure 1A). This helical region had previously been proposed based on secondary structure prediction (Sandu *et al*., 2010), and the AF3 model confirms it as a high-confidence structural element. The predicted Local Distance Difference Test (pLDDT) scores along the α-helix exceed 90, and the PAE (Predicted Aligned Error) plot shows minimal positional error along the helix, suggesting a well-defined and stable structure (Figure 1A).

**Figure 1.**
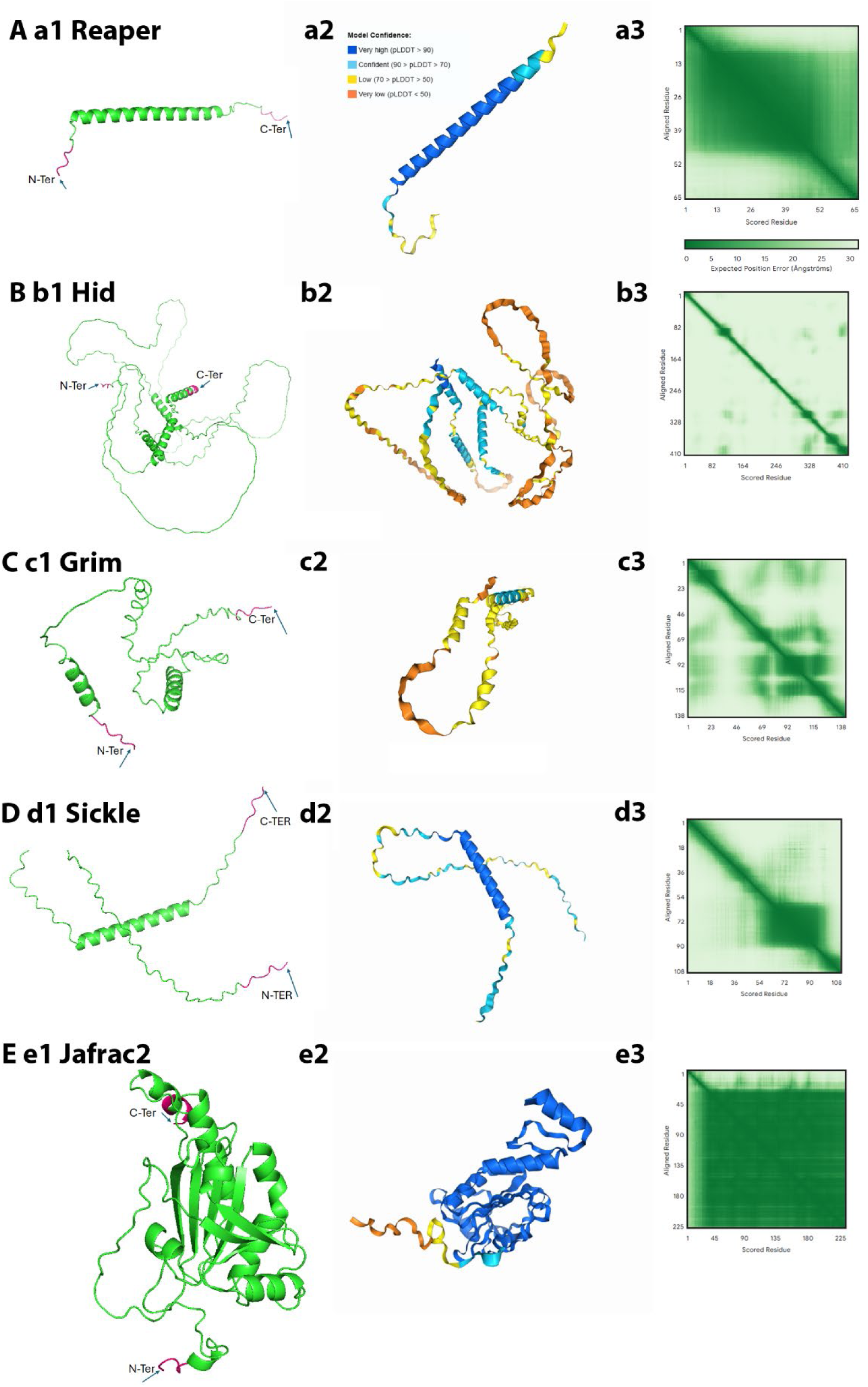
Predicted full-length structures of *Drosophila* IAP antagonists. (**A-E**) AlphaFold3-predicted structures of Reaper (Rpr) (**a1**), Hid (**b1**), Grim (**c1**), Sickle (**d1**), and Jafrac2 (**e1**). (**a2-e2**) Structured regions (primarily α-helices) are colored according to pLDDT (predicted Local Distance Difference Test) confidence values (blue >90, light blue 70-90, yellow 50-70, orange <50), while unstructured regions such as the N-terminal IBM motifs appear as flexible loops. (**a3-e3**) The PAE (Predicted Aligned Error) plots are 2D heatmaps that indicate the positional error (in Ångströms) the model expects when placing one residue of a protein relative to another after optimal alignment. Dark green indicates low positional error, while light green/white indicates high positional error with uncertain relative orientation (such as in flexible linkers or intrinsically disordered regions). Jafrac2 is distinct from the IAP antagonists by forming a compact peroxiredoxin fold with α-helical and β-sheet elements.

Notably, the N-terminal IBM and the C-terminus are unstructured.

Hid is substantially larger (410 aa) (Grether *et al*., 1995), with four predicted α-helices, including the C-terminal TA helix (Figure 1B). These elements exhibit high pLDDT scores, but the inter-helical regions are predicted with low confidence (pLDDT < 50), consistent with extensive intrinsic disorder or structural flexibility. The PAE map reveals widespread uncertainty across the protein, likely due to its low sequence conservation and disordered nature (Figure 1B). As in Rpr, the N-terminal IBM is predicted to be unstructured.

Grim (138 aa) (Chen *et al*., 1996) is predicted to form two α-helices separated by a large, disordered region (Figure 1C). Both the N-terminal IBM and the C-terminal regions are unstructured, and low pLDDT and PAE scores indicate low confidence throughout much of the model.

Sickle (108 aa) (Srinivasula *et al*., 2002; Wing *et al*., 2002) forms a single, well-defined α-helix in the C-terminal half of the protein, with pLDDT values >90 and a consistent PAE map (Figure 1D). The N-terminal IBM is again unstructured.

Mature Jafrac2 (Tenev *et al*., 2002), modeled from aa 18-243, adopts a compact and confidently folded structure (Figure 1E). The predicted peroxiredoxin domain contains both α-helices and β-sheets, yielding high-confidence pLDDT scores (>90). The N-terminal IBM remains unstructured (Figure 1E), in line with the other IAP antagonists.

Altogether, the structural models of these five IAP antagonists reveal a shared architectural theme: short segments of high confidence - usually α-helices - embedded in extensive intrinsically disordered regions (Figure 1). The unstructured regions may confer conformational flexibility, which may facilitate protein-protein interactions. Notably, the N-terminal IBMs of all proteins are predicted to be disordered. This structural flexibility is possibly a functional requirement for binding to the BIR domains of DIAP1. Jafrac2 stands out due to its well-folded peroxiredoxin domain, although its IBM is disordered. These structural predictions provide a foundation for understanding how IAP antagonists execute their pro-apoptotic functions.

### 3.2 Structural predictions of IAP antagonist homodimers

Building on the monomeric structures, we next explored the potential for dimer formation among the five IAP antagonists using AF3. Dimerization has functional significance for at least Rpr and Grim, for which dimerization is known to be both necessary and sufficient for its apoptotic activity (Sandu *et al*., 2010; Yeh and Bratton, 2013). Whether other IAP antagonists form dimers *in vivo* remains unknown.

AF3 predicts that Rpr forms a highly stable homodimer through a parallel alignment of its central α-helical domains (Figure 2A a1). The pLDDT scores in this core region exceed 80, indicating high model confidence, whereas the N- and C-terminal regions including the IBM motifs are disordered and poorly resolved. The PAE map confirms minimal inter-subunit deviation which supports a stable and defined dimer interface (Figure 2A a2). The dimerization surface is mediated by the GH3 motif (aa 32-46), which provides extensive van der Waals contacts between the two subunits with 48 inter-chain atom–atom contacts at ≤ 4.0 Å as well as two salt bridges (Fig. 2A a3). Key GH3 residues (Trp32, Leu35, Val38, Gln45, Tyr46) are among the contact points, in agreement with previous findings by Sandu et al. (2010) (Figure 2A a4).

**Figure 2.**
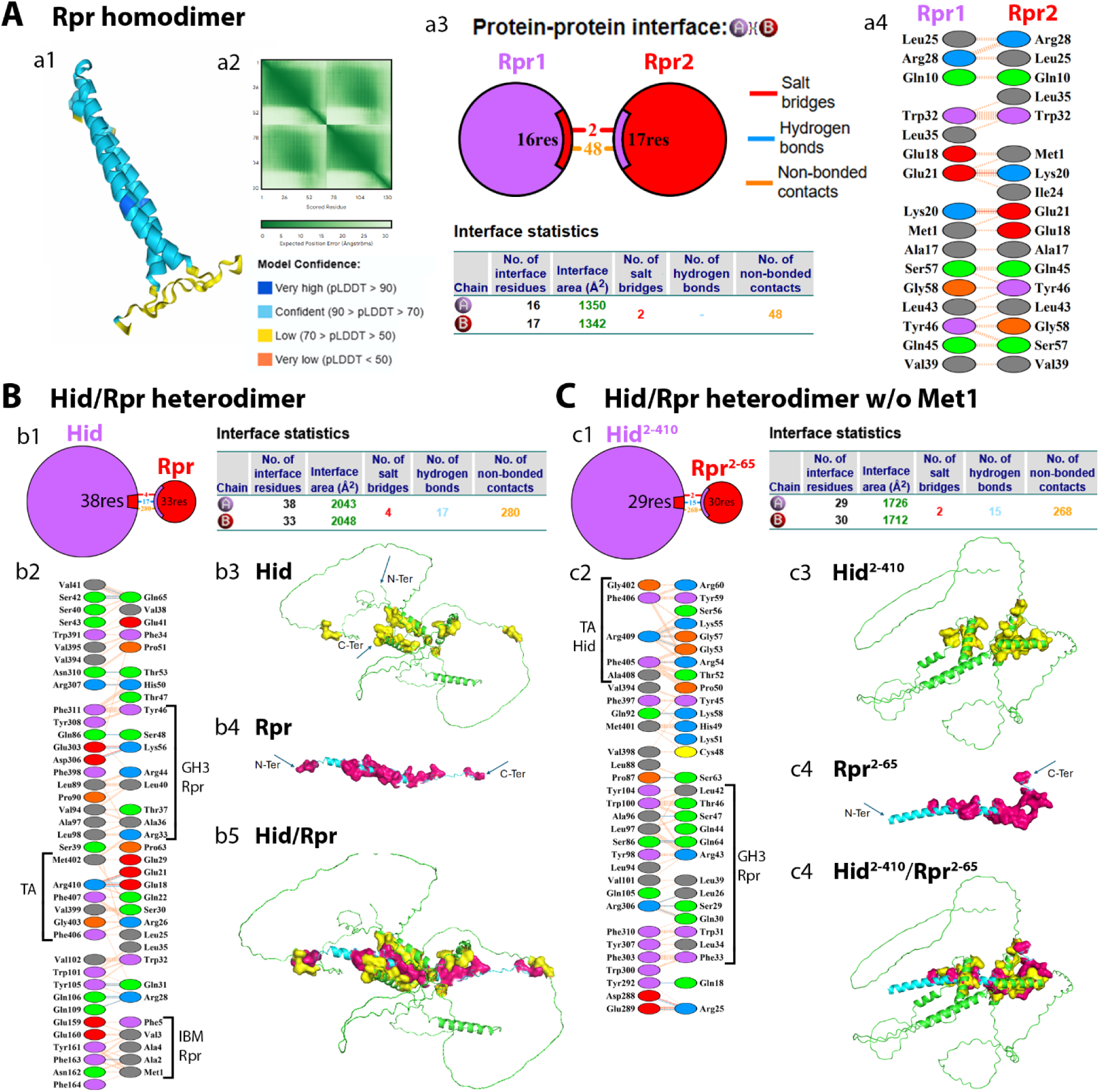
Homo- and heterodimerization of IAP antagonists. (A) (**a1**) Rpr homodimer forms a parallel α-helical coiled-coil interface, primarily mediated by the GH3 motif. The interface includes high-confidence residues (pLDDT >90). (**a2**) The PAE map indicates high positional confidence. (**a3**) Illustration of the protein-protein interface and interface statistics as obtained by PDBsum. Interacting protein chains (Rpr1, Rpr2) are shown as colored circles (purple, red) and are connected by color-coded lines according to the interaction type indicated in the key. The circle size corresponds to the overall surface area of the protein, while the interface is highlighted by a black sector, the span of which reflects the relative interface surface area. (**a4**) Specific residue interactions at the protein-protein interface. The type of interaction is color-coded according to the key in (a3). Hydrogen bonds and salt bridges between amino acids are shown as solid lines. Non-bonded contacts such as van der Waals and aromatic interactions are depicted with striped lines, whose thickness scales with the number of contacts. Non-bonded interactions were considered only if atoms were within 4 Å of each other. Specific atomic details about these interactions are shown in Supplemental Table S1. (B) **Heterodimer of full-length Hid and Rpr.** (**b1, b2**) Protein-protein interface, interface statistics and residue interactions of the heterodimer are as explained in (a3) and (a4). Brackets in (b2) indicate residues of the TA helix of Hid, and the GH3 and IBM motifs of Rpr that are involved in the interaction. (**b3-b5**) PyMOL generated structures of the Hid/Rpr complex. (b3) and (b4) illustrate the structures of the individual proteins (Hid in green, Rpr in cyan; the interacting residues of Hid are highlighted in yellow, the ones of Rpr in red). (b5) depicts the Hid/Rpr complex. N- and C-termini are indicated. These predictions were obtained by ClusPro. (C) **Heterodimer of Hid^2-410^ and Rpr^2-65^ depleted of Met1. (c1, c2).** Protein-protein interface, interface statistics and residue interactions of the heterodimer are as explained in (a3) and (a4). Met1 deletion eliminates the participation of the IBM motif of Rpr in the interaction, while the TA helix of Hid and GH3 motif of Rpr (brackets) remain engaged. (**c3-c5**) PyMOL generated structures of the Hid^2-410^/Rpr^2-65^ complex. (b3) and (b4) illustrate the structures of the individual proteins (Hid in green, Rpr in cyan; the interacting residues of Hid are highlighted in yellow, the ones of Rpr in red). (b5) depicts the Hid^2-410^/Rpr^2-65^ complex. The interaction between the IBM of Rpr with Hid is lost due to Met1 deletion. These predictions were obtained by ClusPro.

Six symmetry-matched residue pairs (Gln10-Gln10, Ala17-Ala17, Arg28-Arg28, Trp32-Trp32, Va39-Val39, and Leu43-Leu43) contact each other across the interface, consistent with the parallel, symmetric alignment of the two Rpr subunits (Fig. 2A a4). The IBM motifs remain solvent-exposed.

In contrast, Hid forms a poorly resolved and unstable dimer with low overall pLDDT scores and a highly variable PAE map (Suppl. Fig. S1A a1). The predicted dimer forms in a parallel configuration, involving interactions between the mid- and C-terminal regions, including the TA helices (Suppl. Fig. S1A a2,a3). However, the low structural confidence across the model calls into question the physiological relevance of this dimer. Notably, the TA helices which do interact in the dimer are proposed to associate with the outer mitochondrial membrane (OMM).

The Grim homodimer exhibits low pLDDT scores across most of the structure (Suppl. Fig. S1B b1). Interestingly, AF3 predicts an anti-parallel alignment between the two α-helices in the mid-region of the protein (Suppl. Fig. S1B b2,b3), in contrast to the parallel arrangement observed in Rpr and Hid dimers. The overall low confidence suggests a flexible or transient interaction.

For Sickle, AF3 predicts a relatively stable parallel homodimer (Suppl. Fig. S1C), with high pLDDT values in the α-helical core and slightly reduced confidence at the interface.

Dimerization involves residues in the mid- and C-terminal regions, with a minor and low-confidence contribution from the IBM.

Jafrac2 forms an anti-parallel homodimer involving residues near both termini (Suppl. Fig. S1D). While the central peroxiredoxin domain is well-folded and contributes to dimer stability, the IBM and terminal regions remain disordered. The dimer is more stable than those of Hid and Grim, as indicated by higher pLDDT scores in the core (Suppl. Fig. S1D).

In summary, Rpr forms a robust and well-supported homodimer, with high-confidence scores across the dimer interface and clear alignment with functional data. In contrast, the predicted homodimers of Hid, Grim, Sickle, and Jafrac2 are characterized by low structural confidence or high flexibility. Given these limitations, only the Rpr homodimer will be later further examined for its apoptotic function. Notably also, all homodimers retain disordered N-terminal IBMs, consistent with their role in flexible, transient interactions with DIAP1.

### 3.3 Heterodimeric interactions of IAP antagonists and implications for apoptotic complex formation

Previous studies have shown that Rpr, Hid, and Grim localize to mitochondria during the apoptotic process (Claveria *et al*., 2002; Haining et al., 1999; Olson *et al*., 2003). Rpr and Grim contain a conserved central GH3 motif, which is essential for mitochondrial localization, although GH3 itself is not a classical mitochondrial targeting signal. In contrast, the C-terminal TA helix of Hid integrates it into the outer mitochondrial membrane (OMM). Sandu et al. (2010) provided biochemical evidence that Rpr can associate with Hid and that this interaction facilitates the localization of Rpr to the OMM, a key requirement for its apoptotic activity.

To gain structural insight into these heterotypic interactions, we used AF3 to model heterodimeric complexes of Rpr, Hid, Grim, and Sickle. Among them, the Rpr/Hid interaction is the most biologically relevant and best supported by prior functional studies.

AF3 predicts that Rpr and Hid form a stable heterodimer with a well-defined and extensive interface (Figure 2B b1,b2). Notably, the interface is dominated by hydrophobic residues, forming a nonpolar core stabilized by discrete polar interactions. The computed lowest binding energy for this complex is -1395 kcal/mol, consistent with a stable complex.

Interestingly, Rpr retains its rod-like conformation, while Hid adapts its conformation to form extensive interactions with Rpr (Figure 2B b3,b4). This conformational plasticity may enable Hid to engage multiple binding partners. The GH3 motif of Rpr (aa 32-46), which is known to mediate interaction with Hid (Sandu *et al*., 2010), engages residues within α-helix 1 (aa 90-110) and α-helix 2 (aa 302-320) of Hid (Figure 2B b2,b5). In addition, the IBM of Rpr interacts with a disordered region of Hid (aa 159-165), and the mid-region (aa 18-33) of Rpr contacts the C-terminal TA helix of Hid. This latter interaction may facilitate the recruitment of Rpr to the OMM by anchoring it via the TA domain of Hid.

From this AF3 prediction, a particularly intriguing and novel insight arises from the role of the initiating methionine (Met1) in regulating the Rpr/Hid interaction. It is well-established that Met1 must be cleaved from Rpr, Hid, and Grim to expose the IBM for binding to the BIR domains of DIAP1 (Wu *et al*., 2000). However, our structural model shows that in the Rpr/Hid complex, the IBM, in particular Met1, of Rpr engages in multiple interactions with residues 161-164 of Hid (Figure 2B b2). This suggests that in the Rpr/Hid complex, the IBM domain of Rpr may be masked by Hid, making this protein interaction potentially anti-apoptotic in nature.

However, when Met1 is removed from Rpr, the IBM of Rpr is now released (Figure 2C), but the overall stability of the complex drops significantly (lowest binding energy changes from -1395 to -1284.6 kcal/mol).

This finding reveals an unexpected and previously unrecognized dual role for Met1 of Rpr which introduces a potential checkpoint in apoptosis. While Met1 must be removed to permit binding of the IBM to DIAP1 and trigger apoptosis, its presence is important for stabilizing the Rpr/Hid complex. This tradeoff suggests a finely tuned regulatory mechanism in which Met1 removal not only licenses apoptotic activity, but also destabilizes pro-apoptotic complexes. These insights provide a new layer of understanding into the spatial and temporal control of apoptosis in *Drosophila* (see Discussion).

We also modeled other heterodimers, including Hid/Grim, Rpr/Grim, and Hid/Sickle (Suppl. Fig. S2A–C). Among these, the Hid/Grim complex displayed the most extensive interface, involving 52 residues from Hid and 64 from Grim, forming 30 hydrogen bonds, 7 salt bridges, and 399 van der Waals contacts, with a lowest binding energy of -2122 kcal/mol (Suppl. Fig. S2A). Unlike the linear Rpr, Grim has a more compact, globular structure, and Hid adopts a distinct conformation to accommodate this partner, which highlights its structural plasticity. (Suppl. Fig. S2A a3,a4). The GH3 motif of Grim (aa 85-101) interacts with both α-helix 1 and the TA helix of Hid. The IBM of Grim is only partially engaged, with residues 3-7 forming limited contacts.

The Rpr/Grim heterodimer is weaker (lowest binding energy: -1274 kcal/mol), with interactions primarily between the α-helical and C-terminal regions of Rpr and the C-terminal half of Grim (Suppl. Fig. S2B). The GH3 motifs of both proteins are involved, but their IBMs are not. The conformation of Grim in this complex closely resembles its structure in the Grim/Hid dimer, despite differences in the interacting residues.

The Hid/Sickle complex (lowest binding energy: -1656 kcal/mol) involves 36 residues from Hid and 42 from Sickle (Suppl. Fig. S2C). The GH3 motif of Sickle interacts with α-helix 2 of Hid, but again, the IBMs of both proteins are not involved. Hid adopts yet another conformation, further demonstrating its role as a structurally adaptable scaffold.

Together, these models highlight both known and novel features of IAP antagonist interactions and the adaptability of Hid to bind partners with distinct shapes. The Rpr/Hid complex stands out as the most functionally relevant and structurally robust heterodimer, whereas other complexes - though intriguing - remain to be validated *in vivo*.

### 3.4 Structural prediction of full-length DIAP1 reveals a compact, globular domain organization

DIAP1 is composed of two BIR domains and one C-terminal RING domain (Figure 3A).

**Figure 3.**
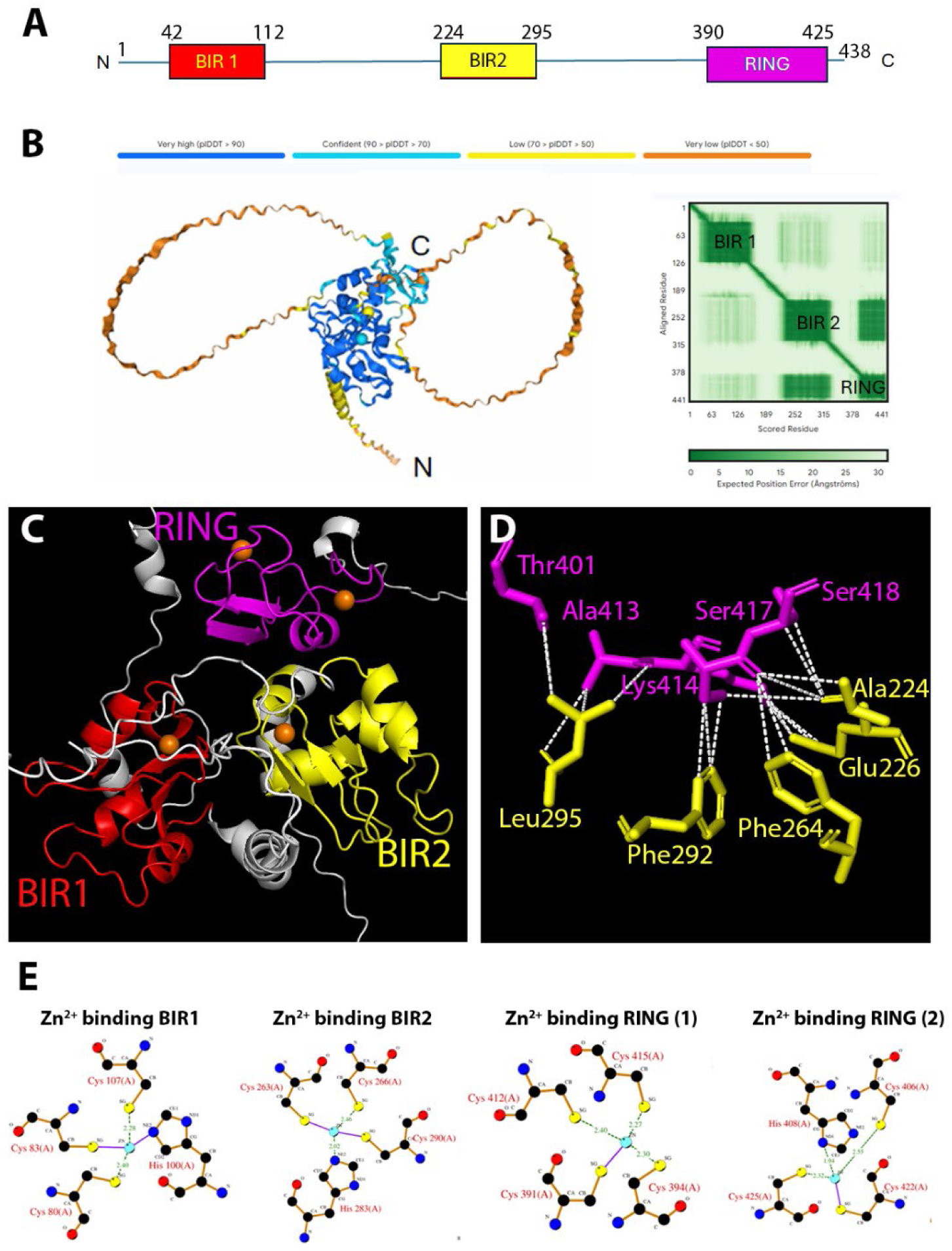
Structural model of full-length DIAP1. (A) Domain organization of full-length DIAP1. The colors for the domains (BIR1 red; RIR2 yellow and RING magenta) are consistently used in this paper. (B) AF3 prediction of full-length DIAP1 reveals a compact globular arrangement of the BIR1, BIR2, and RING domains, with unstructured loops connecting them. Coloring reflects pLDDT confidence according to the key. The PAE map indicates high global confidence within the BIR1, BIR2 and RING domains, and indicates that the BIR2 and RING domains form a structurally stable module. (C) PyMOL-generated high magnification view of the arrangements of the BIR1, BIR2 and Ring domains. Color-coding is according to (A). Linker regions are in white. The Zn^2+^ ions are indicated by orange spheres. (D) Detailed interaction view of the BIR2/RING interaction. Salt bridges and non-bonded contacts (van der Waals interactions) between residues of the BIR2 and RING domains are shown as white dashed lines. Only residues that are involved in the interaction are shown and labeled. BIR2 residues are in yellow, RING residues in magenta. See Suppl. Table S1 for specific details of these interactions. (E) Zn^2+^ coordination in the BIR and RING domains involves canonical histidine and cysteine residues.

Structural knowledge to date has focused on individual domains, which have been resolved at high resolution by X-ray crystallography. These studies have provided invaluable insight into how the IBMs of IAP antagonists interact with the isolated BIR domains (Chai *et al*., 2003; Wu *et al*., 2000; Wu *et al*., 2001; Yan *et al*., 2004). However, it has remained unknown how these domains are arranged within the intact protein, specifically, whether they are connected by flexible linkers in an extended "beads-on-a-string" arrangement, or whether they adopt a more compact, folded architecture.

To address this gap, we used AF3 to predict the 3D structure of full-length DIAP1, including chelated Zn² ions. Our model reveals a globular, compact fold in which the BIR1, BIR2, and RING domains are not randomly oriented, but are tightly packed into a cohesive structural core (Figure 3B). Each domain is predicted with very high confidence (pLDDT > 90), while the linker regions connecting them are characterized by lower confidence scores and predicted to form flexible, unstructured loops. These findings are further supported by the PAE map, which shows low positional uncertainty within individual domains and a strong spatial correlation between BIR2 and the RING, while increased variability is observed in the interdomain regions (Figure 3B). This arrangement suggests that DIAP1 functions as an integrated structural unit rather than a modular scaffold.

Closer inspection of the DIAP1 model reveals that BIR1 and BIR2 are positioned on opposite surfaces of the core (Figure 3C), potentially allowing simultaneous or cooperative engagement of IAP antagonists via their IBM motifs. In contrast, the RING domain is oriented adjacent to the BIR domains (Figure 3C), and exposes a distinct surface patch that likely corresponds to the binding site of the E2 ubiquitin-conjugating enzyme. Importantly, detailed analysis shows that BIR2 and the RING domain are interconnected by a network of close contacts, including two salt bridges and numerous van der Waals interactions (Figure 3D; for a detailed list of the interactions at atomic level see Suppl. Table S1). These electrostatic and hydrophobic interactions provide strong evidence that BIR2 and the RING form a structurally stable module which is also reflected by the PAE map (Figure 3B). This arrangement suggests a degree of functional compartmentalization, in which substrate recognition (via the BIR domains) and ubiquitin transfer (via the RING domain) are structurally separated, but held within a unified scaffold. These insights provide a structural basis for understanding how DIAP1 integrates caspase inhibition, binding of IAP antagonists and ubiquitin ligase functions within a spatially coordinated molecular platform.

The AF3 model also accurately recapitulates Zn² coordination within each domain, consistent with experimental observations. Specifically, BIR1 and BIR2 each coordinate a single Zn^2+^ ion through conserved histidine and cysteine residues, while the RING domain coordinates two Zn^2+^ ions required for structural integrity and catalytic activity (Figure 3E). This correct placement of metal-binding residues further supports the reliability of the predicted domain architecture.

In summary, the structural prediction of full-length DIAP1 provides the first comprehensive view of its domain organization. Rather than behaving as a flexible, modular protein, DIAP1 adopts a compact, globular fold with tightly packed BIR and RING domains. This architecture likely facilitates coordinated regulation of caspase inhibition and IAP antagonist binding.

### 3.5 Differential interactions of IAP antagonists with DIAP1

DIAP1 is an essential inhibitor of apoptosis in *Drosophila*, functioning to suppress caspase activation in most somatic cells (Goyal *et al*., 2000; Lisi *et al*., 2000; Wang *et al*., 1999). In apoptotic cells, DIAP1 is antagonized by pro-apoptotic IAP antagonists (Reaper, Hid, Grim, Sickle, and Jafrac2), which bind to DIAP1 through their N-terminal IBM motifs, thereby triggering caspase activation and cell death (Goyal *et al*., 2000; Vucic et al., 1997; Vucic *et al*., 1998; Yan *et al*., 2004).

Early structural studies using X-ray crystallography provided key insights into these interactions by resolving complexes between IBM-containing 10-mer peptides (residues 2–11, lacking the initiator Met) of Rpr, Hid, and Grim and the isolated BIR domains of DIAP1 (Wu *et al*., 2001; Yan *et al*., 2004). These studies showed that the IBM peptides of Rpr, Hid and Grim can bind to both BIR1 and BIR2 of DIAP1. Further biochemical binding studies demonstrated the Rpr and Grim can interact evenly with both isolated BIR1 and BIR2 domains, while Hid, Sickle and Jafrac2 have a binding preference for BIR2 (Zachariou et al., 2003). However, due to technical challenges in expressing and crystallizing full-length, multi-domain proteins, these studies did not address how the full-length IAP antagonists interact with full-length DIAP1.

Thus, it remained unclear whether additional interactions outside the IBM contribute to DIAP1 binding, and whether IBM binding influences domain arrangements in full-length DIAP1. To address these questions, we used AF3 to systematically model and compare the interactions of IBM 10-mer peptides with isolated BIR domains of DIAP1 (validating previous work), IBM peptides with full-length DIAP1, and full-length IAP antagonists (lacking N-terminal Met1) with full-length DIAP1.

#### 3.5.1 Validation of known IBM peptide/BIR interactions

First, we used AF3 to replicate the known structures of IBM peptides (aa 2-11) from Rpr, Hid, and Grim in complex with isolated BIR2 domains (Wu *et al*., 2001). Table 2 presents the residues of BIR2 that interact with the IBM motifs of Rpr, Hid, and Grim. A group of five residues of BIR2 (Gly269, Met271, Asp277, Gln282, Trp286) forms hydrogen bonds (highlighted in green in Table 2) with residues in all IBMs (Ala1, Val/Ile2, Phe/Tyr4, Tyr/Phe5), while several additional BIR2 residues (up to 11) form non-bonded contacts with the IBMs (highlighted in yellow in Table 2; see also Suppl. Fig. S3; for a detailed list of the interactions at atomic level see Suppl. Table S1). These data align perfectly with the crystallographic binding studies of Wu et al. (2001) (Table 2), validating the AF3 approach. We also extended this analysis to Sickle and Jafrac2, and observed similar but weaker interactions with BIR2 compared to Rpr, Hid or Grim (Table 2; Suppl. Table S1; Suppl. Fig. S3).

**Table 1:** Alignment of the IBM motives of the IAP Antagonists Amino acid sequences of the first 10 residues, highlighting the key difference in position 3.

| IAP-antagonists | IBM motif (2-11) |
| --- | --- |
| Reaper | A V **A** F Y I P D Q A |
| Hid | A V **P** F Y L P E G G |
| Grim | A I **A** Y F I P D Q A |
| Sickle | A I **P** F F E E E H A |
| Jafrac2 (18-27 aa) | A K **P** E D N E S C Y |

**Table 2:**
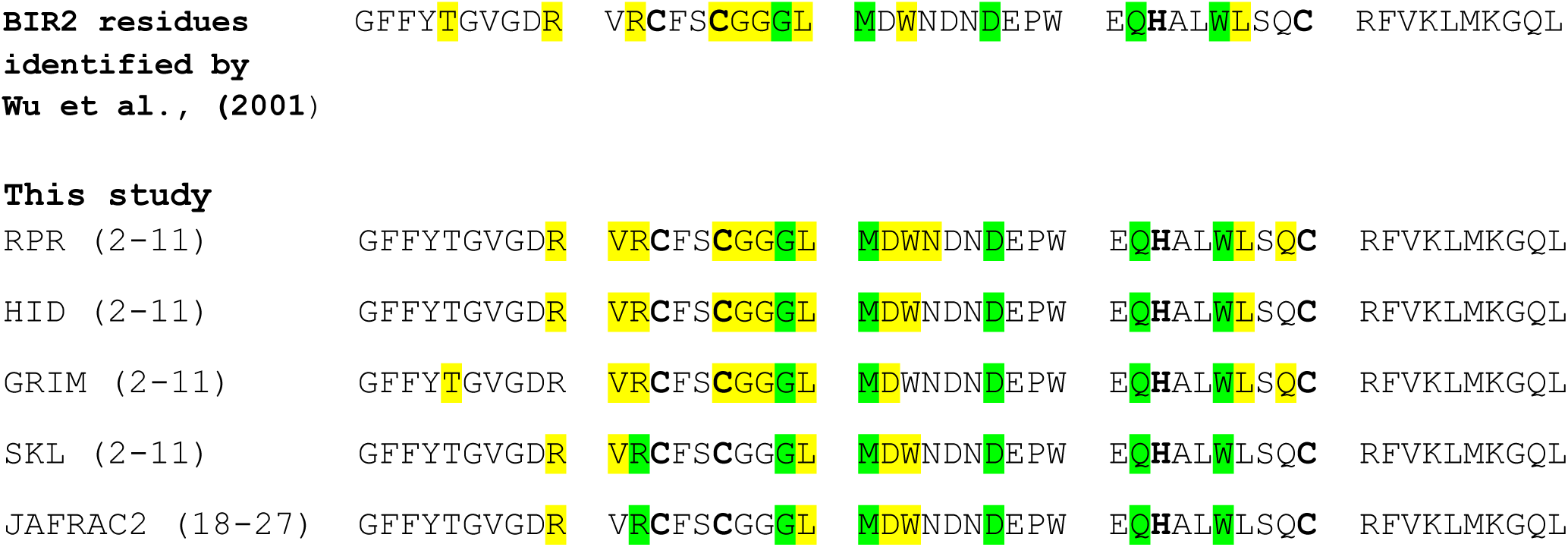
Interaction Table Between Isolated BIR2 domain and IBM Peptides (2-11 Amino Acids) (see also Supplementary Figure S3 and Supplementary Table S1) Aligned are residues 251-300 of DIAP1 (encompassing BIR2). Residues forming H-bonds and non-bonded contacts with the IBM peptides (aa 2-11) of the indicated IAP antagonists are highlighted in green (common residues are Gly269, Met271, Asp277, Gln282, Trp282); those residues making non-bonded contacts are highlighted in yellow. Cys and His residues in bold are chelating Zn^2+^ ions. The data predicted by AF3 match very well the data obtained by Wu et al. (2001) obtained for the Hid and Grim IBM peptides.

#### 3.5.2 IBM peptides with full-length DIAP1

Next, we modeled binding of the IBM peptides with full-length DIAP1. Interestingly, when having the choice between BIR1 and BIR2, all IBM peptides preferentially interacted with BIR2 (Suppl. Fig. S4; Table 3; Suppl. Table S1), including the IBM of Rpr, which previously showed preferred specificity for BIR1 in domain-only contexts (Tenev et al., 2005; Zachariou *et al*., 2003). Despite variation in IBM conformation (Suppl. Fig. S4), the same conserved BIR2 residues (Gly269, Met271, Asp277, Gln282, Trp286) consistently formed hydrogen bonds with residues 1, 2, 4, and 5 of the IBMs (Table 3). This suggests that within the structural context of full-length DIAP1, BIR2 may be more accessible or favorably oriented for IBM engagement.

**Table 3:**
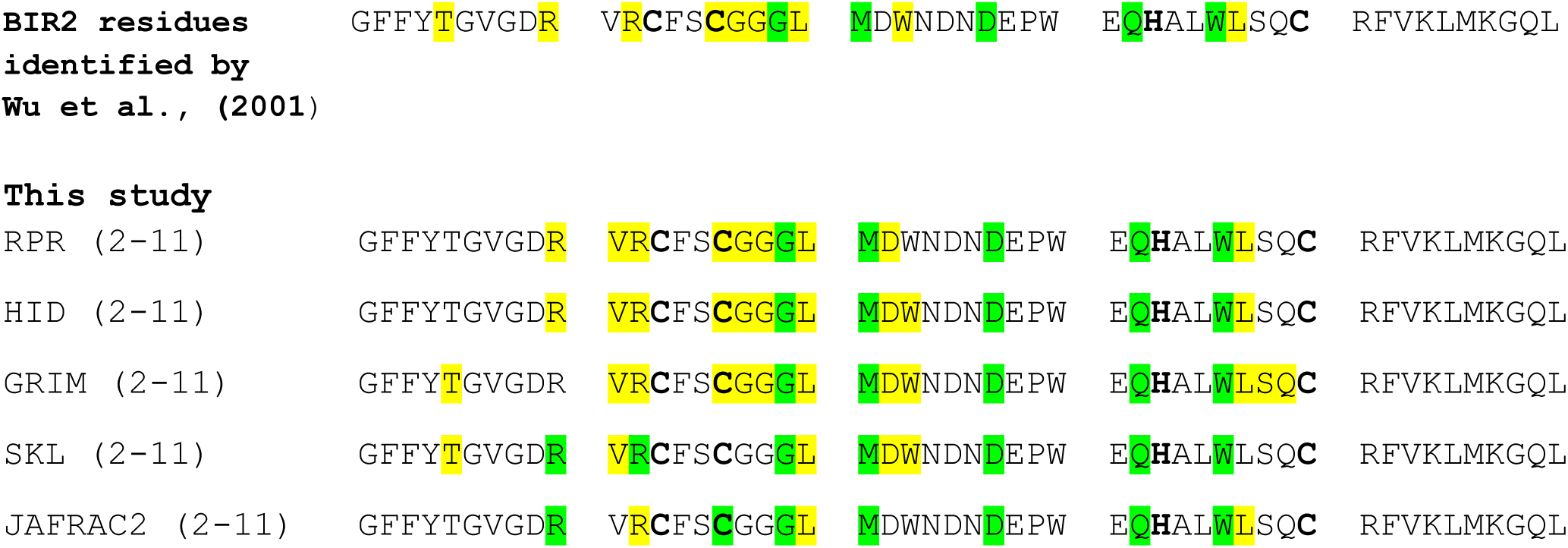
Interaction Table Between Full-Length DIAP1 and IBM Peptides (2-11 Amino Acids) (see also Supplementary Figure S4 and Supplementary Table S1) In these AF3 predictions, the IBM peptides of all IAP antagonists interact with the BIR2 domain, not with the BIR1 domain. Aligned are residues 251-300 of DIAP1 (encompassing BIR2). Residues forming H-bonds and non-bonded contacts with the IBM peptides (aa 2-11) of the indicated IAP antagonists are highlighted in green (common residues are Gly269, Met271, Asp277, Gln282, Trp282); those residues making only non-bonded contacts are highlighted in yellow. Cys and His residues in bold are complexed by Zn^2+^ ions. The data predicted by AF3 match very well the data obtained by Wu et al. (2001) which were obtained for the Hid and Grim IBM peptides in complex with the isolated BIR2 domain.

#### 3.5.3 Full-length IAP antagonists with full-length DIAP1

Modeling full-length IAP antagonists (all lacking the N-terminal Met1) with full-length DIAP1 revealed striking differences in domain engagement and uncovered novel, context-dependent determinants of binding specificity beyond the canonical IBM/BIR interface (Figure 4).

**Figure 4.**
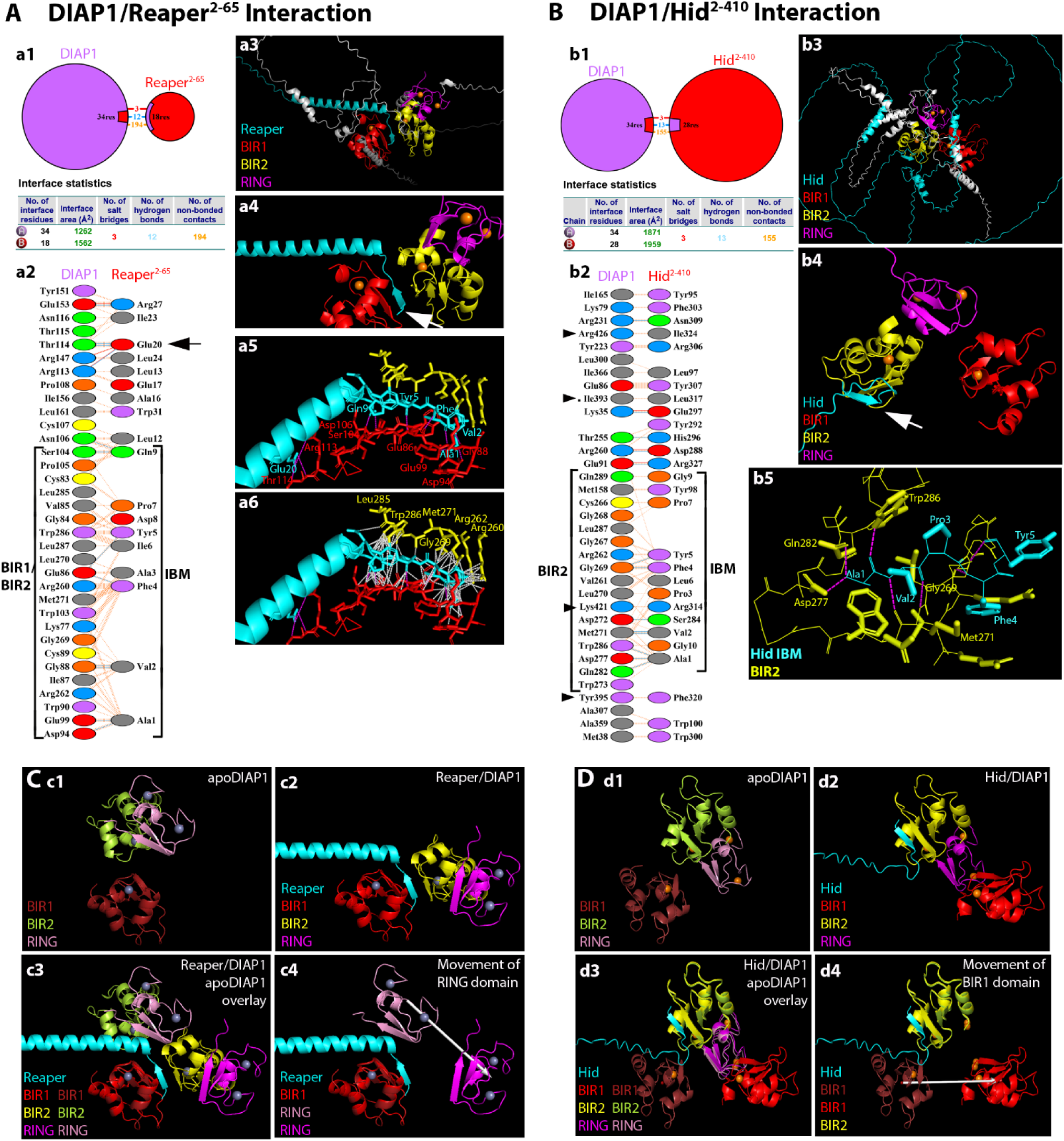
Interactions of full-length DIAP1 with Rpr^2-65^ and Hid^2-410^ as predicted by AF3. (A) **The DIAP1/Rpr^2-65^ complex**. (**a1**) Illustration of the protein-protein interface and interface statistics as obtained by PDBsum. (**a2**) Specific residue interactions between the protein-protein interface. Rpr^2–65^ binds primarily to BIR1 via its IBM, but also forms non-bonded contacts with BIR2 (brackets). Additional residues from the central α-helix, in particular Glu20 (black arrow), contribute to the interaction with DIAP1. (**a3**) Overall view of the DIAP1/ Rpr^2-65^ complex. Rpr^2-65^ is shown in cyan, BIR1 in red, BIR2 yellow and the RING in magenta. Zn^2+^ ions are orange spheres. (**a4**) Higher magnification view of the DIAP1/Rpr^2-65^ complex. The IBM of Rpr inserts mainly into the groove of the BIR1 domain (white arrow), but is also in proximity to BIR2. (**a5**) Detailed interaction view of the IBM/BIR1 interaction. Only H-bonds (magenta dashed lines) are shown to demonstrate their exclusive formation between BIR1 and the IBM. All residues from both the BIR1 and IBM that participate in H-bond formation are labeled in red and cyan, respectively. Also emphasized is Glu20 from Rpr^2-65^, which establishes strong interactions with DIAP1 residues Arg113 and Thr114 through H-bonds, salt bridges, and non-bonded contacts (represented solely by magenta H-bond lines). (**a6**) Detailed interaction view of the IBM/BIR1 and IBM/BIR2 interaction. Identical view as (a5), but non-bonded contacts are also included as white dashed lines. The IBM of Rpr^2-65^ mainly interacts with BIR1, but also makes non-bonded contacts with BIR2. Key residues of the BIR2 domain involved in non-bonded contacts are labeled in yellow. Residues of BIR1 and the IBM of Rpr are not labeled for clarity (but see a5). (B) **The DIAP1/Hid^2-410^ complex**. (**b1**) Illustration of the protein-protein interface and interface statistics as obtained by PDBsum. (**b2**) Specific residue interactions between the protein-protein interface. The IBM of Hid^2–410^ binds to BIR2 (brackets). Hid also uses residues of its α-helix 2 (aa 302-320) to contact the RING domain of DIAP1 (arrowheads). (**b3**) Overall view of the DIAP1/Hid^2-410^ complex. (**b4**) Higher magnification view of the DIAP1/Hid^2-410^ complex. The IBM of Hid (cyan) inserts into the groove of BIR2 (yellow) (white arrow). (**b5**) Detailed interaction view of the IBM/BIR2 interaction. Critical residues for this interaction are labeled. Purple dashed lines indicate H-bonds. Non-bonded contacts are omitted for clarity. (C) **Conformational changes of DIAP1 upon Rpr^2-65^ binding.** Structural overlay of unbound (apo) DIAP1 and the DIAP1/Rpr^2-65^ complex aligned on the BIR1 domain. (**c1**) Only the BIR1 (ruby), BIR2 (limegreen) and RING (pink) domains of apoDIAP1 are shown. Linker regions are hidden for clarity. (**c2**) Only the BIR1 (red), BIR2 (yellow) and RING (magenta) domains of DIAP1 in complex with Rpr^2-65^ (cyan) are shown. (**c3**) Using BIR1 as anchor, the structural overlay of apoDIAP1 and complexed DIAP1 reveals a striking conformational rearrangement of the BIR2 and RING domains. (**c4**) A white arrow indicates the movement of the RING domain in response to Rpr^2-65^ binding. The RING domain moves by ∼27 Å and pivots by ∼58° toward BIR1, bringing it ∼11 Å closer to BIR1. (D) **Conformational changes of DIAP1 upon Hid^2-410^ binding.** Structural overlay of apoDIAP1 and the DIAP1/Hid^2-410^ complex aligned on the BIR2 domain. (**d1, d2**) Color code of BIR1, BIR2 and RING of apoDiap1 and complexed DIAP1 as in (c1,c2). Hid IBM in cyan. Linker regions are hidden for clarity. (**d3**) In the BIR2-anchored frame, the RING domain remains essentially unchanged relative to BIR2. In contrast, the BIR1 domain undergoes a pronounced relocation. (**d4**) A white arrow indicates the movement of the BIR1 domain in response to Hid^2-410^ binding, shifting its centroid by ∼55 Å and rotating by ∼138° relative to apoDIAP1.

As a full-length protein, Rpr^2-65^ (Met1 deleted) binds strongly to BIR1 via its IBM (Figure 4A; Table 4A; Suppl. Table S1). This interface is stabilized by multiple hydrogen bonds and non-bonded contacts between key BIR1 residues (Glu86, Gly88, Asp94, Glu99, Ser104, Asn106) and the side chains or main-chain atoms of Ala1, Val2, Phe4, Tyr5, and Gln9 within the IBM of Rpr (Figure 4A a2,a4,a5; Table 4A,C; Suppl. Table S1). These specific interactions firmly anchor the IBM in the BIR1 pocket. Interestingly, there are also weaker non-bonded contacts between the IBM and BIR2, such that the IBM is positioned between the two BIR domains but engaged asymmetrically, i.e. strongly anchored to BIR1, while only loosely contacting BIR2 (Figure 4A a2,a6; Table 4B; Suppl. Table S1). Beyond the IBM of Rpr, additional interactions between the central α-helical region of Rpr and residues between BIR1 and BIR2 appear to stabilize this conformation and steer the IBM toward BIR1, overriding its default affinity for BIR2 as a peptide (as seen in the last section) (Figure 4A a2,a5; Suppl. Table S1). This highlights a critical role for non-IBM regions in modulating their binding specificity to DIAP1. The RING domain of DIAP1 is not directly involved in this interaction.

**Table 4:**
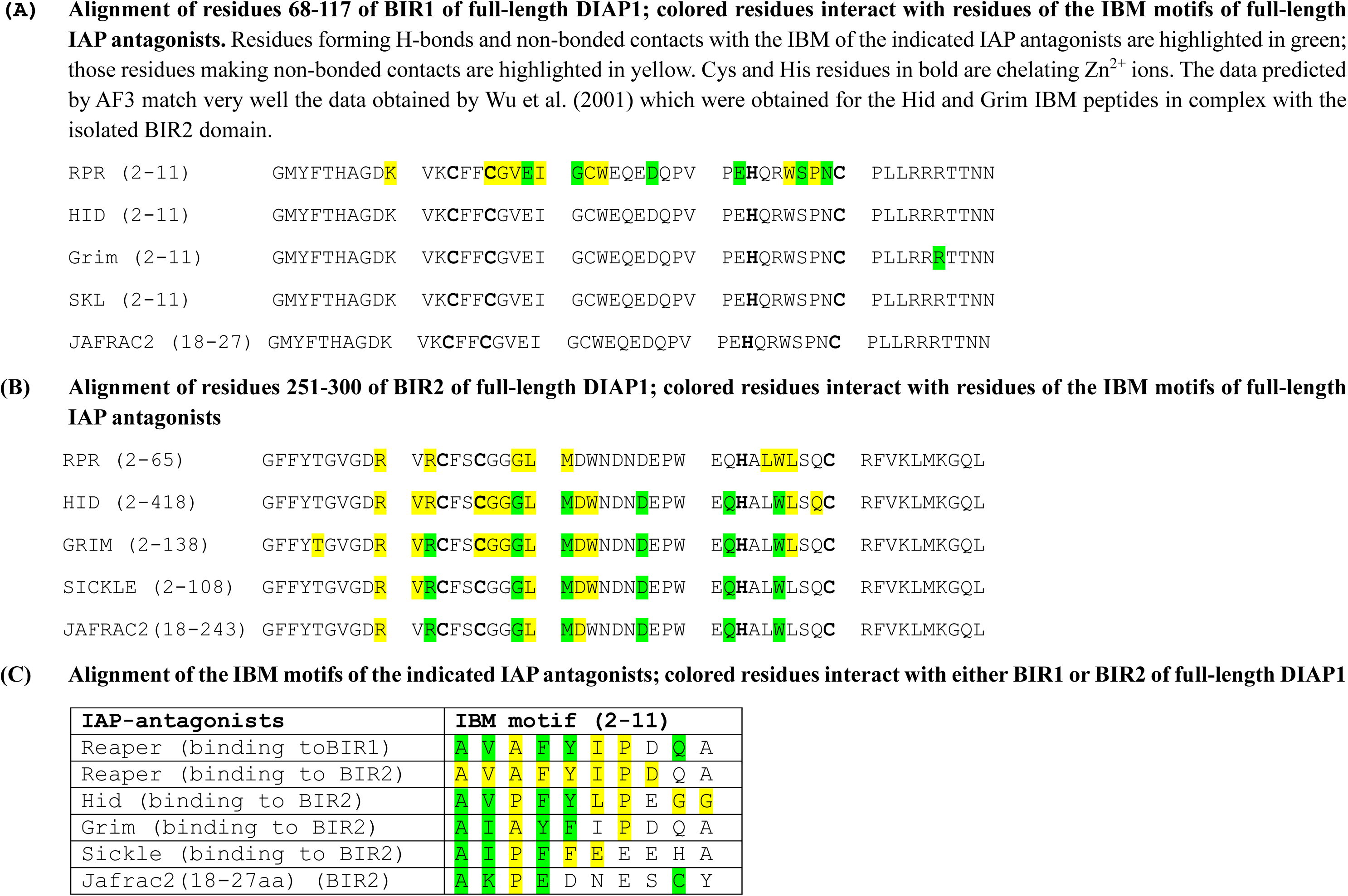
Interaction of BIR1 and BIR2 of full-length DIAP1 with the IBM motifs of the full-length IAP antagonists. (see also Figure 4, Supplementary Figure S5, and Supplementary Table S1)

Comparison of unbound (apo)DIAP1 with DIAP1 in complex with Rpr revealed striking conformational rearrangements of the BIR2 and RING domains when structures were aligned on the invariant BIR1 domain (Figure 4C). Upon Rpr binding, the BIR2 domain undergoes a large spatial rearrangement, shifting its centroid by ∼31 Å relative to apoDIAP1 and rotating by 55–60° around its axis with respect to BIR1 (Figure 4C c3). The RING domain, tracked via the Zn^2+^ centroid, is likewise repositioned, moving by 27 Å and pivoting by 58° toward BIR1 (Figure 4C c3,c4) demonstrating that Rpr binding induces a hinge-like motion across the BIR1-BIR2-RING module which brings the RING domain ∼11 Å closer to BIR1. Functionally, such repositioning of the RING domain might be highly significant. In the apo state, the RING lies distal from BIR1, consistent with its ability to ubiquitylate caspases. By contrast, in the Rpr-bound state, the RING is redirected toward the BIR1/IBM interface, a geometry that would facilitate DIAP1 self-ubiquitylation. These findings support the model that Reaper not only displaces caspases from DIAP1, but also actively remodels the architecture of DIAP1 to favor its autoubiquitylation and degradation.

In the DIAP1/Hid^2-410^ complex, the IBM of Hid^2–410^ engages exclusively with BIR2 of DIAP1 (Figure 4B b2,b4,b5; Table 4B; Suppl. Table S1), docking into the canonical groove where BIR2 envelops the IBM from two sides, which effectively occludes access by BIR1. The IBM contribution is mediated by Ala1, Val2, Phe4, and Tyr5 of Hid, which bury against a BIR2 wall centered on Gly269, Leu270, Met271, and Trp286, with auxiliary polar contacts from Arg262, Asp272/277, and Gln282 of DIAP1 (Figure 4B b5; Table 4B,C). Notably, all IBM contacts detected within 4.0 Å are to BIR2. Outside the IBM/BIR2 interface, a secondary contact patch involves the RING domain (Lys392, Ile393, Cys394, Tyr395, Lys414, Cys415, Ser418, Pro423) of DIAP1 which engages with residues in α-helix 2 of Hid (Phe310, Gly313, Arg314, Leu317, Gly322, Ile324). These peripheral contacts likely enhance complex stability and enforce a bipartite binding mode, in which a tight IBM/BIR2 core is complemented by RING-proximal touches that help orient and stabilize the complex.

Alignment of apoDIAP1 with DIAP1 in complex with Hid revealed that the BIR2 domains of the apo and complex structures overlap almost perfectly (Figure 4D d1-d3). In this BIR2-anchored frame, the RING domain (defined by the Zn² midpoint) also remains essentially unchanged relative to BIR2, consistent with the idea that BIR2 and RING form a structurally stable module (see also Figure 3D). By contrast, the BIR1 domain undergoes a pronounced relocation, shifting its centroid by ∼55 Å and rotating by 138° relative to apoDIAP1 (Figure 4D d3,d4) demonstrating that Hid binding induces a dramatic displacement of BIR1 away from the BIR2-RING core, while the BIR2-RING arrangement itself is unchanged.

Functionally, this contrasts with the effect of Rpr binding to DIAP1. Whereas Rpr draws the RING toward BIR1 to promote DIAP1 self-ubiquitylation, Hid appears to stabilize the BIR2-RING unit and dislodge BIR1, thereby disrupting caspase engagement without repositioning the RING. Importantly, however, this observation does not necessarily mean that Hid cannot induce DIAP1 autoubiquitylation. α-helix 2 of Hid contributes additional contacts with the RING of DIAP1 (Figure 4B b2). Such direct helix/RING interactions could alter the local environment of the RING and redirect its ubiquitin ligase specificity toward autoubiquitylation, albeit through a distinct mechanism from the conformational repositioning triggered by Rpr.

Grim^2-138^ engages BIR2 via its IBM, with no detectable interaction with BIR1 (Suppl. Fig. S5A; Table 4; Suppl. Table S1). Outside the IBM, only seven additional residues from Grim contact DIAP1, mostly near the BIR2 and RING regions. The overall interaction appears weaker and more limited in scope than the Rpr or Hid complexes.

Skl^2-138^ binds DIAP1 through canonical IBM/BIR2 interactions, with no BIR1 engagement (Table 4; Suppl. Fig. S5B; Suppl. Table S1). Notably, additional residues in the C-terminal region of Sickle (Gln82, His87, Ser90, Lys97, Thr99) form interactions with the RING domain of DIAP1, suggesting a more extended interaction interface than seen with Grim.

These results demonstrate that full-length context significantly influences the specificity and stability of IAP antagonist/DIAP1 complexes. While IBMs alone preferentially target BIR2, the presence of additional structured elements in the full-length proteins can redirect or reinforce binding to a particular BIR domain. This will be specifically addressed in the case of Rpr in the next section.

### 3.6 Domain swapping and mutational analysis reveal IBM and backbone determinants of BIR-binding specificity of Rpr

The findings from the previous section raise an intriguing question: Why does full-length Rpr uniquely bind to BIR1 (and to some extent to BIR2) of DIAP1, while the Rpr IBM peptide alone and all other IAP antagonists - both as peptides and full-length proteins - preferentially bind to BIR2? This question is even more intriguing given the sequence similarities of the IBMs of the IAP antagonists (Table 1). To address this question, we designed a series of domain swaps, in which we replaced the IBM of one IAP antagonist with that of another, and of point mutations using AF3 modeling. These *in silico* constructs allowed us to dissect the determinants of BIR-domain specificity in the context of full-length Rpr.

First, we exchanged the IBM of Rpr with that of Grim which usually interacts with BIR2 (Suppl. Figure S5A), generating a chimera termed G/Rpr^2-65^. However, when docked with full-length DIAP1, G/Rpr^2-65^ maintained strong interaction with BIR1, suggesting that the Rpr backbone can guide even a heterologous IBM toward BIR1 (Figure 5A; Suppl. Figure S6A; Suppl. Table S1). Conversely, the reciprocal chimera R/Grim^2-138^ bound preferentially to BIR2, suggesting that the IBM of Rpr does not have an intrinsic affinity for BIR1 (Figure 5B; Suppl. Figure S6B; Suppl. Table S1). These IBM swapping experiments imply that backbone residues of Rpr play an instructive role for positioning its IBM for productive interaction with BIR1.

**Figure 5.**
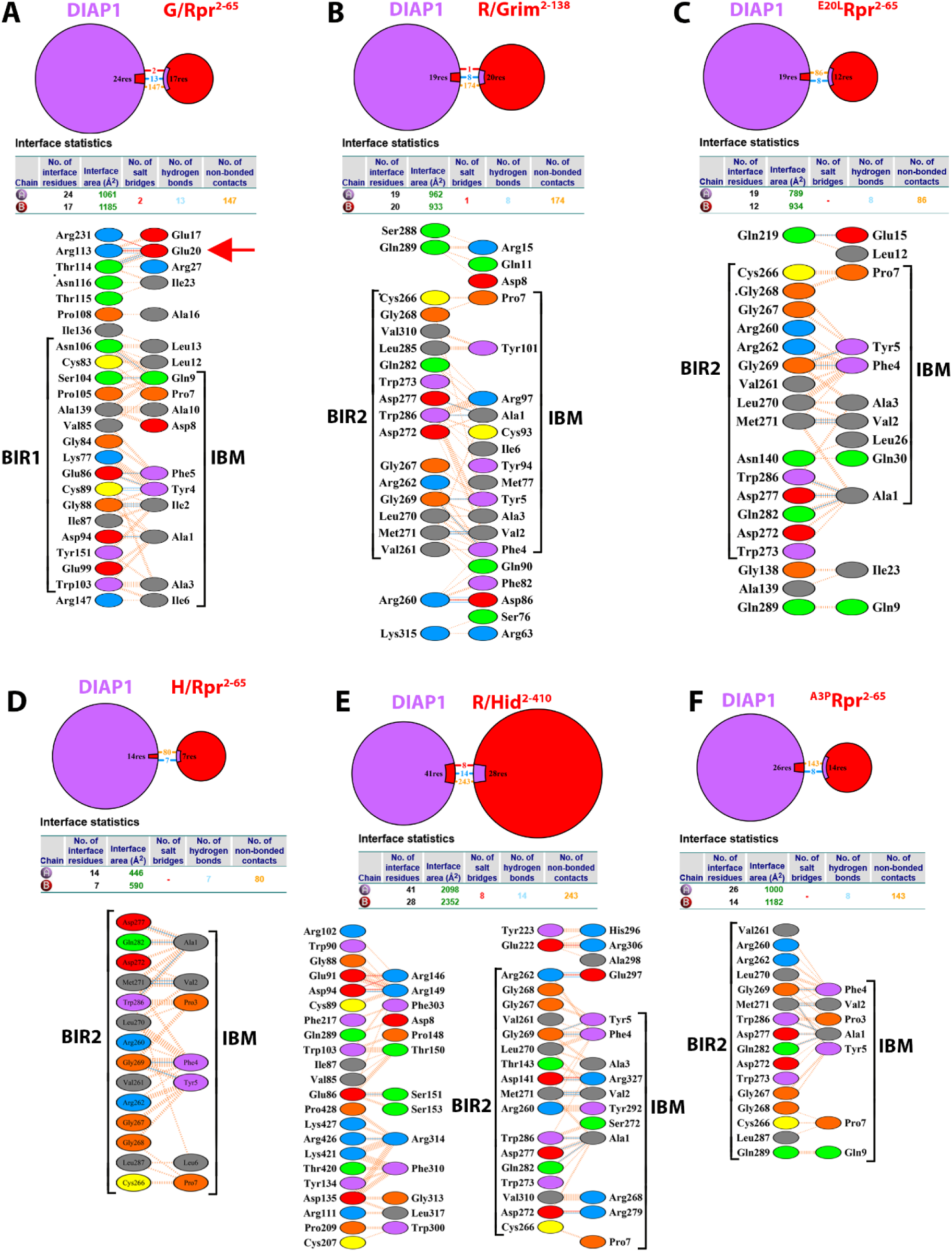
Domain swap and point mutation analysis of IBM binding specificity. All panels show protein-protein interfaces, interface statistics and residue interactions of the heterodimers obtained by PDBsum according to Figure 2A a3 and a4. The brackets indicate residues of the BIR domains of Diap1 and the IBM motifs of the chimeric and mutated IAP antagonists that are involved in the interaction. PyMOL-generated images of the protein complexes are shown in Supplementary Figure S6. (A) G/Rpr^2–65^ (Grim IBM in Rpr backbone) binds BIR1. (B) R/Grim^2–138^ binds BIR2. (C) Substitution of Glu20 with Leu in the backbone of Rpr^2-65^ (^E20L^Rpr^2-65^) redirects binding from BIR1 to BIR2. (D) H/Rpr^2–65^ and (**E**) R/Hid^2–410^ both bind BIR2, showing IBM-determined specificity. (**F**) Substitution of Ala3 with Pro in the IBM of Rpr^2-65^ (^A3P^Rpr^2-65^) switches binding specificity from BIR1 to BIR2.

Further analysis of the Rpr/DIAP1 interaction interface revealed six backbone residues of Rpr (Leu12, Leu13, Glu17, Glu20, Ile23, Arg27) that establish stable contacts with DIAP1 (Figure 4A a2; Figure 5A; Suppl. Table S1). Among these, Glu20 is of particular interest because it establishes multiple hydrogen bonds and salt bridges with Arg113 and Thr114 of DIAP1, located immediately adjacent to BIR1 (Figure 4A a2,a5; Figure 5A; Suppl. Table S1). This unusually strong interaction appears to bias the orientation of the Rpr IBM toward BIR1. To directly test this hypothesis, we substituted Glu20 with Leu (^E20L^Rpr^2-65^) and modeled the mutant protein in complex with DIAP1 using AF3. Strikingly, this single substitution redirected the IBM preference from BIR1 to BIR2 (Figure 5C; Suppl. Figure S6C; Suppl. Table S1). This result highlights Glu20 as a molecular pivot that dictates IBM positioning and provides a critical molecular understanding of how the Rpr backbone may influence the specificity of DIAP1 binding.

However, the influence of the IBM sequence for BIR binding preference cannot be discounted. When we replaced the Rpr IBM with that of Hid (producing H/Rpr^2-65^), the resulting chimera now bound to BIR2, not BIR1 (Figure 5D; Suppl. Figure S6D; Suppl. Table S1). The reverse chimera, R/Hid^2-410^, also bound exclusively to BIR2 (Figure 5E; Suppl. Figure S6E; Suppl. Table S1). These results suggest that in some cases, specific IBM residues can override the influence of the protein backbone.

A key difference between the IBMs of Rpr and Hid lies at position 3: Rpr contains Ala3, while Hid contains Pro3 (Table 1). Because such a substitution could critically influence how the IBM engages with different BIR domains, we tested whether replacing Ala3 with Pro (^A3P^Rpr^2-65^) alters the binding specificity of Rpr. Strikingly, this single substitution switched the binding preference of Rpr from BIR1 to BIR2 (Figure 5F; Suppl. Figure S6F; Suppl. Table S1).

Structural modeling revealed the likely basis for this effect. In the Rpr/BIR1 complex, Ala3 packs closely against Trp103 of BIR1, forming stabilizing hydrophobic contacts (Figure 4A a2; Suppl. Table S1). However, a Pro at position 3 introduces steric hindrance with Trp103, preventing efficient binding. In contrast, Pro3 fits well into the BIR2 binding pocket, interacting favorably with Gly269, Leu270, and Trp286 (Figure 4B b2; Suppl. Table S1), which are core IBM-interacting residues in BIR2.

These results reveal a layered mechanism of specificity in IAP antagonist/DIAP1 interactions. While all IBMs display an intrinsic affinity for BIR2, backbone residues in Rpr, specifically Glu20, can redirect the IBM toward BIR1. However, the presence of Pro at position 3 acts as a dominant determinant, overriding backbone influence and re-directing specificity to BIR2. This demonstrates the sensitive interplay between local side-chain chemistry and overall protein architecture. Interestingly, also, while the H/Rpr^2-65^ chimera and the ^A3P^Rpr^2-65^ mutant successfully redirect Rpr to BIR2, neither construct establishes interactions with DIAP1 outside the IBM motif (Figure 5D,F). This suggests that Rpr lacks evolutionary adaptation for BIR2 binding.

### 3.7 Structural insights into the interaction between Jafrac2^18-242^ and DIAP1

The thioredoxin peroxidase Jafrac2, encoded by *Prx4*, is a structurally and functionally unusual IAP antagonist in *Drosophila*. In contrast to canonical IAP antagonists such as Rpr, Hid, and Grim which localize to mitochondria and are dedicated to promoting apoptosis, Jafrac2 resides in the ER under normal physiological conditions, where it functions as an oxidoreductase that promotes disulfide bond formation during protein folding (Tenev *et al*., 2002). During its ER import, the N-terminal signal peptide of Jafrac2 is cleaved off, generating mature Jafrac2^18-242^ and exposing an IBM at the new N-terminus (Tenev *et al*., 2002). Upon apoptotic stimuli, Jafrac2^18-242^ is rapidly released into the cytosol, where it can act as an IAP antagonist, thereby promoting caspase activation and cell death (Tenev *et al*., 2002). This functional duality distinguishes Jafrac2 from the mitochondrial IAP antagonists, which lack alternative cellular roles and are dedicated solely to apoptosis. Mutations that disrupt Jafrac2^18-242^/DIAP1 binding suppress its pro-apoptotic activity, underscoring the importance of this interaction (Tenev *et al*., 2002).

Jafrac2^18–242^ protein was confidently predicted by AF3, with the exception of the unstructured IBM at the N-terminus, which remains flexible (Figure 1E). To better understand how Jafrac2^18–242^ functions as an IAP antagonist, we used AF3 to model its interaction with full-length DIAP1. As expected, based on its IBM sequence, which contains a Proline at position 3, the IBM of Jafrac2^18-242^ engages exclusively with BIR2 of DIAP1 (Figure 6A-D; Suppl. Table S1). Key residues in the IBM (Ala18, Lys19, Pro20) were central to this interaction. Ala18 formed hydrogen bonds with Gln282, Asp277, and Tyr286; Lys19 interacted with Asp272 and Met271; and Pro20 contributed van der Waals contacts with Tyr286 (Figure 6B; Supplementary Table S1). These interactions closely parallel the IBM-mediated contacts observed in other BIR2-binding IAP antagonists.

**Figure 6.**
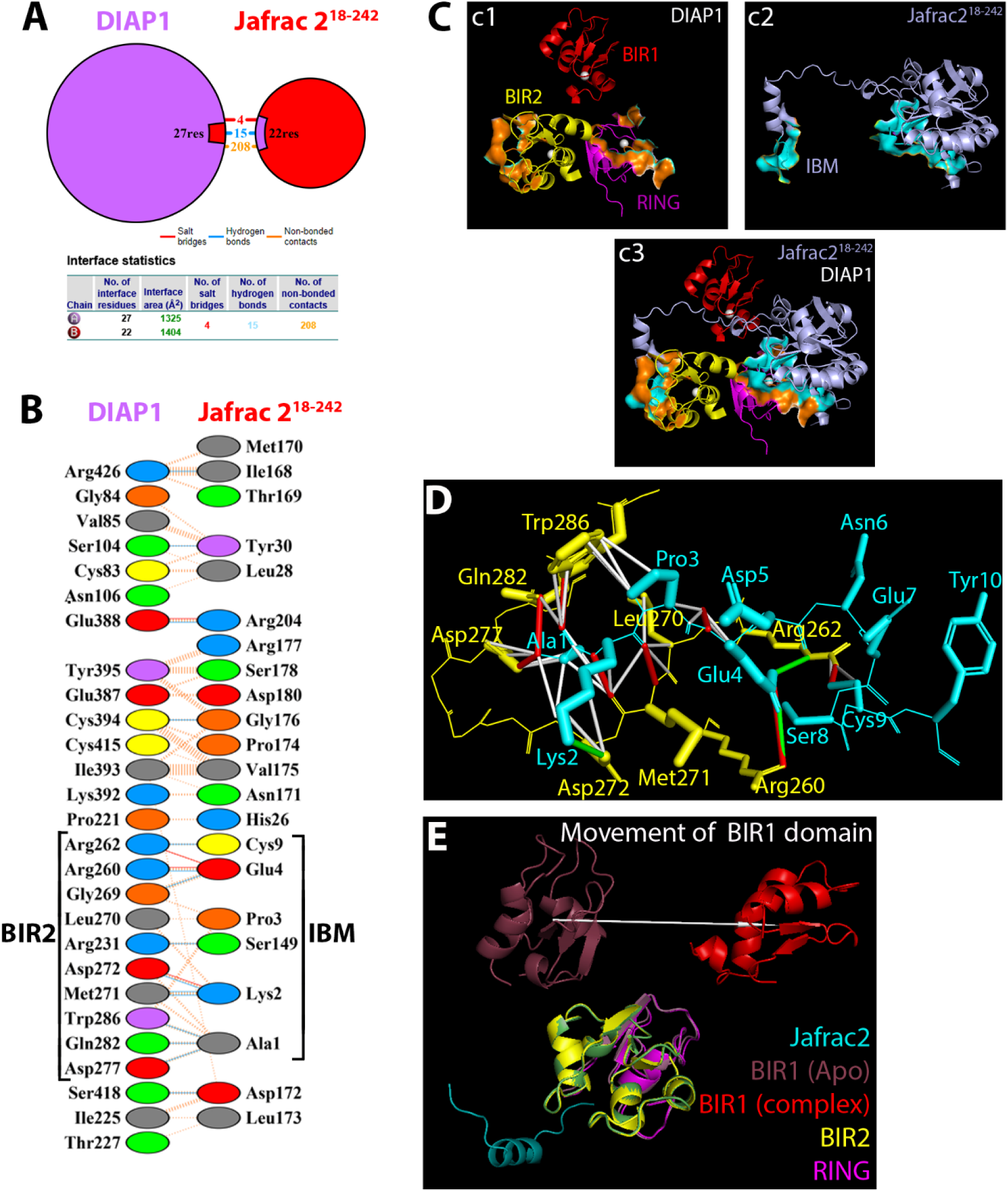
Jafrac2^18-242^ engages a canonical interaction with the BIR2 of DIAP1. (A) Illustration of the protein-protein interface and interface statistics as obtained by PDBsum. (B) Specific residue interactions at the protein-protein interface. The IBM of Jafrac^18–242^ binds to BIR2 of DIAP1 (brackets). Jafrac^18–242^ also uses residues outside the IBM to interact with the RING domain of DIAP1. (C) PyMOL-generated view of the DIAP1/Jafrac^18–242^ complex. (**c1**) and (**c2**) illustrate the structures of the individual proteins. The domains of DIAP1 are color-coded according to Figure 3A. The interacting regions in the BIR2 and RING of DIAP1 are highlighted in orange in surface mode. The corresponding interacting regions in Jafrac2^18-242^ are shown in cyan in surface mode. (**c3**) depicts the DIAP1/Jafrac2^18-242^ complex. (D) Detailed mechanistic view of the IBM/BIR2 interaction. Critical residues for this interaction are labeled. Red lines indicate H-bonds, green lines are salt bridges and white lines are non-bonded contacts. (E) Conformational changes of DIAP1 upon Jafrac^18-242^ binding. In the BIR2-anchored frame, the RING domain remains essentially unchanged relative to BIR2. In contrast, there is significant movement of the BIR1 domain in response to Jafrac2^18-242^ binding, shifting its centroid by ∼41 Å and rotating by ∼77° relative to apoDIAP1 (white arrow).

Notably, Jafrac2^18–242^ also engages DIAP1 outside the IBM/BIR2 interface. Structural modeling reveals that residues from the peroxiredoxin core domain of Jafrac2^18–242^ interact with regions of BIR1 and the RING domain of DIAP1 (Figure 6B,C; Suppl. Table S1). These extended contacts likely contribute to complex stabilization.

Comparison of apoDIAP1 with the Jafrac2-bound structure shows that the BIR2 regions align closely, indicating that this domain is not detectably altered by Jafrac2 binding (Figure 6E). Within this BIR2-anchored frame, the RING domain also superimposes well, underscoring the notion that BIR2 and RING behave as a rigid structural module. In contrast, Jafrac2 binding causes BIR1 to shift by ∼41 Å and rotate by ∼77° (Figure 6E, white arrow). This demonstrates that Jafrac2 selectively mobilizes BIR1 without perturbing the BIR2-RING module. A qualitatively similar movement is observed with Hid binding (Figure 4D), but in this case BIR1 undergoes an even more pronounced relocation (∼55 Å and ∼138°) indicating that while both IAP antagonists drive BIR1 away from its apo position, the extent of this displacement is greater for Hid.

Nevertheless, these structural predictions confirm and extend previous findings, establishing Jafrac2^18–242^ as a *bona fide* IAP antagonist that uses its N-terminal IBM to engage BIR2, while also using its folded core for additional interactions with DIAP1. This work provides the structural framework for understanding how a redox enzyme evolved a secondary pro-apoptotic function through domain modularity and subcellular re-localization.

### 3.8 Oligomeric state-dependent inhibition of DIAP1 by Rpr and Hid

Previous studies have demonstrated that dimerization of Reaper is necessary and sufficient for its pro-apoptotic activity (Sandu *et al*., 2010). Additionally, Rpr and Hid have been shown to form higher-order structural complexes that are crucial for apoptotic activity (Sandu *et al*., 2010). These findings collectively establish oligomerization as a functionally relevant feature for apoptosis. Building on this, we used AF3 to model how oligomeric states of Rpr and its interaction with Hid affect DIAP1 binding and inhibition.

#### 3.8.1 Reaper homodimer binding to DIAP1

We first modeled the trimeric complex of DIAP1 (chain A) bound to a Rpr homodimer (chains B and C) (Figure 7A,B). As expected, the IBM of the first Rpr subunit (Rpr1) engages the BIR1 domain of DIAP1, forming hydrogen bonds and salt bridges with canonical binding residues (Gly84, Ile87, Cys89, Asp94, Glu99, Arg102) (Figure 7C,F,G; Suppl. Table S1).

**Figure 7.**
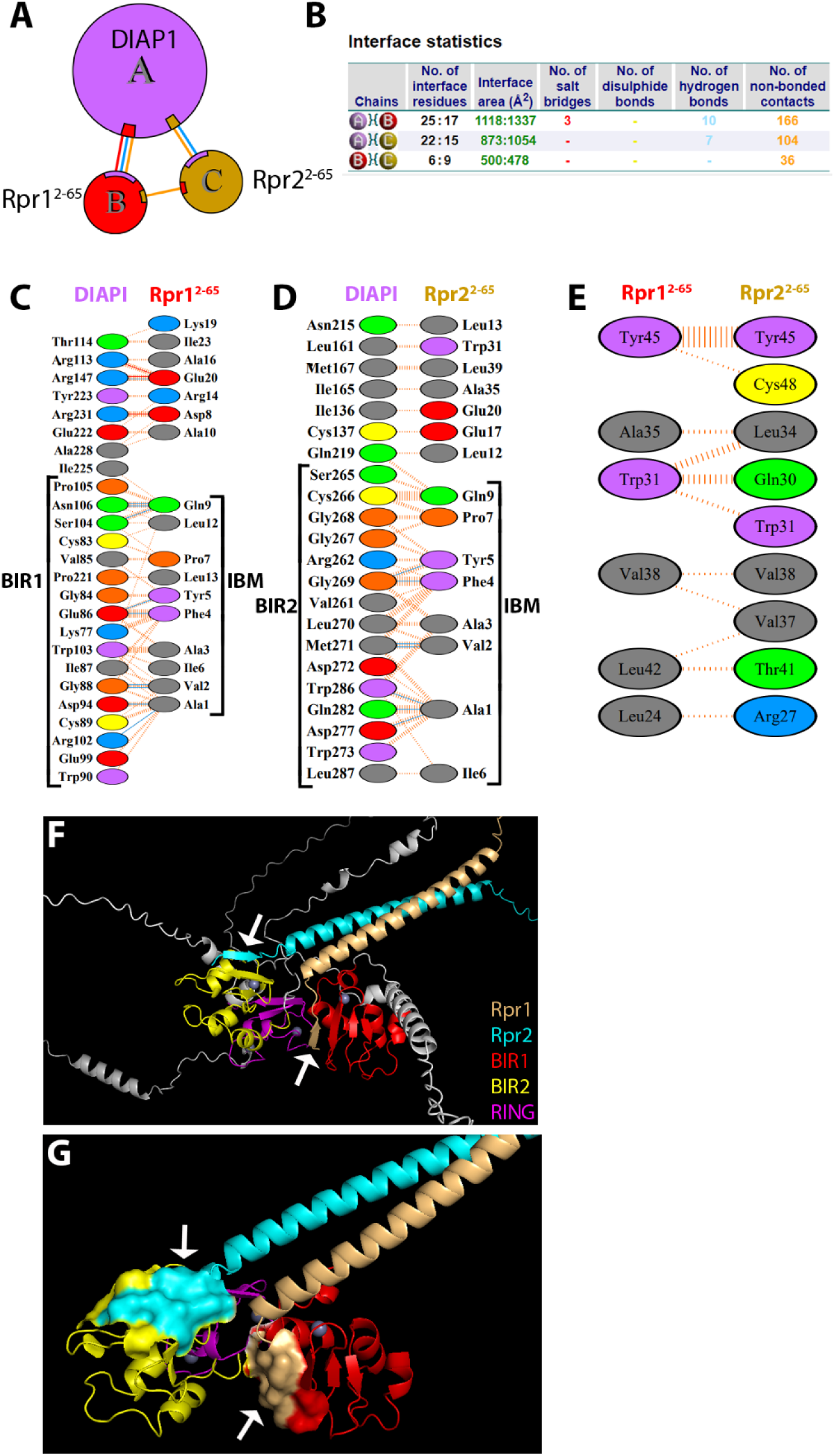
Ternary complex of DIAP1 with a Rpr^2-65^ dimer. (**A, B**) Illustration of the protein-protein interfaces and interface statistics as obtained by PDBsum. (**C-E**) Specific residue interactions at the protein-protein interface as obtained by PDBsum. The IBM of Rpr1 binds BIR1 (C); the IBM of Rpr2 binds BIR2 (D) (indicated by brackets). The Rpr dimer interface weakens upon DIAP1 binding (E). (F) PyMOL-generated overall view of the DIAP1/Rpr1/Rpr2 complex. Rpr1 and Rpr2 are colored in light orange and cyan, respectively. The BIR1, BIR2 and the RING domain of DIAP1 are colored in red, yellow and magenta, respectively. Linker regions are in white. White arrows point to the IBM/BIR1 and IBM/BIR2 interface. (G) Higher magnification view of the DIAP1/Rpr1/Rpr2 complex. The interactions of the IBMs with the BIR domains are indicated in surface mode (white arrows). Linker regions are omitted for clarity.

Interestingly, since BIR1 is occupied by Rpr1, the IBM of the second Rpr subunit (Rpr2) instead interacts with the BIR2 domain, engaging typical residues such as Leu270, Met271, Trp273, Asp277, Gln282, and Tyr286 (Figure 7D,F,G; Suppl. Table S1).

Quantitatively, the Rpr1/BIR1 interaction is stronger, forming 3 salt bridges, 10 hydrogen bonds, and 166 van der Waals contacts (from now on referred to as 3/10/166), compared to the Rpr2/BIR2 interface (0/7/104) (Figure 7B,C). Notably, however, both of these individual interfaces are weaker than the DIAP1/Rpr heterodimer modeled earlier (3/12/191) (Figure 4A). This suggests that DIAP1 binding affinity per Rpr subunit is reduced in the trimeric context,.

Interestingly, the Rpr1/Rpr2 interaction within the DIAP1-bound trimer is also weakened compared to the Rpr/Rpr homodimer. While the Rpr/Rpr homodimer alone is stabilized by (2/0/48) (Figure 2A), the interaction between the two Rpr subunits in the ternary DIAP1/Rpr/Rpr complex is reduced to (0/0/36) (Figure 7E), suggesting that DIAP1 binding alters the stability of the Rpr dimer interface. These findings highlight the dynamic nature of these interactions.

#### 3.8.2 DIAP1 binding by a Rpr dimer and Hid monomer

Next, we modeled the quaternary complex containing DIAP1, Hid and a Reaper homodimer (DIAP1/Hid/Rpr1/Rpr2) (Figure 8A,B). In this quaternary complex, DIAP1 is simultaneously engaged by Hid and the two Rpr subunits, producing a highly cooperative inhibitory assembly. Hid binds extensively to the BIR2 domain of DIAP1 via its N-terminal IBM, which inserts into the conserved hydrophobic groove formed by Gly269, Leu270, and Trp286 (Figure 8C,F,G). In addition to the BIR2/IBM interaction, and similar to the Diap1/Hid heterodimer (Figure 4B), Hid also makes contact with residues of the RING domain through its α-helix 2 (Figure 8C, arrowheads).

**Figure 8.**
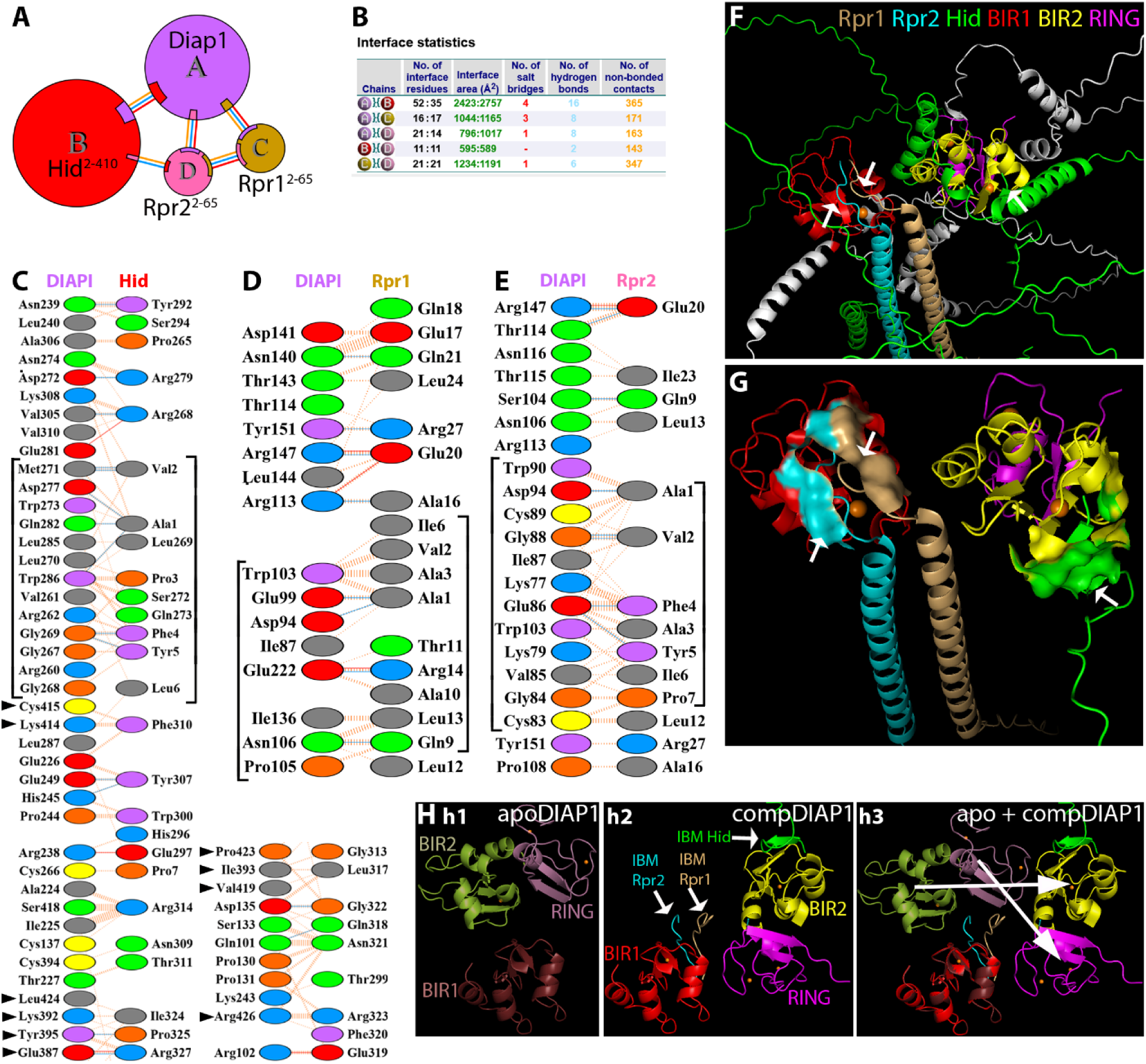
Quaternary complex of DIAP1, Hid^2-410^ and Rpr^2-65^ dimer. (**A, B**) Illustration of the protein-protein interfaces and interface statistics as obtained by PDBsum. (**C-E**) Specific residue interactions at the protein-protein interfaces as obtained by PDBsum. The IBM of Hid binds to BIR2 (C). Arrowheads point to the interaction of α-helix 2 of Hid with residues of the RING domain of DIAP1. The IBMs of both Rpr1 and Rpr2 share binding with BIR1 (D,E) (see brackets). Additional chain/chain interactions are shown in Supplemental Figure S7. (**F**) PyMOL-generated overview of the DIAP1/Hid/Rpr1/Rpr2 complex. Hid, Rpr1 and Rpr2 are colored in green, light orange and cyan, respectively. The BIR1, BIR2 and the RING domain of DIAP1 are colored in red, yellow and magenta, respectively. White arrows point to the IBM/BIR1 and IBM/BIR2 interfaces. (**G**) Higher magnification view of the DIAP1/Hid/Rpr1/Rpr2 complex. The interactions of the IBMs with the BIR domains are indicated in surface mode, indicated by white arrows. Linker regions are omitted for clarity. (**H**) Overlay of apoDIAP1 and DIAP1 in complex with Hid and two Reaper molecules. The structures were aligned using the BIR1 domains as anchors. For clarity, linker regions between the DIAP1 domains have been omitted, and only the IBM (the first 10 amino acids) of Hid and Rpr1/2 are displayed. In the overlay, the BIR2 domain of complexed DIAP1 is displaced by ∼18 Å and rotated by ∼35°, while the RING domain is displaced by ∼25 Å and rotated by ∼55°, relative to apoDIAP1. White arrows in (h2) mark the interfaces where the IBMs engage their respective BIR domains. Large white arrows in (h3) indicate the movement of the BIR2 and RING domains of DIAP1 upon binding of Hid, Rpr1 and Rpr2.

Interestingly, the BIR1 domain of DIAP1 is engaged by the two Rpr subunits simultaneously. Each Rpr subunit contributes its IBM to the limited BIR1 surface, but the engagement is asymmetric. The IBM of Rpr1 appears to make less intense contact with BIR1 than the IBM of Rpr2. Notably, in this quaternary context, the IBMs of both Rpr subunits also contact one another directly (Suppl. Figure S7C) which is absent in the isolated Rpr homodimer and in the DIAP1/Rpr/Rpr trimer (Figures 2A and 7). This additional interaction reinforces the Rpr1/Rpr2 interface and might compensate for the weaker individual BIR1 contacts in the quaternary complex. There is also an interaction between Hid and Rpr2 in the quaternary complex involving α-helix 1 of Hid and the central helix of Rpr2 (Suppl. Figure S7B).

Quantitative comparisons of the interfaces highlight the cooperative architecture of the quaternary DIAP1/Hid/Rpr1/Rpr2 complex. The DIAP1/Hid interaction is much stronger in the quaternary complex compared to the heterodimeric complex of Fig. 4B, expanding from 3 salt bridges, 13 H-bonds and 155 non-bonded contacts (3/13/155) in the heterodimer to (4/16/365) in the quaternary state (Figure 4B, Figure 8B; Suppl. Table S1). Importantly, outside of the canonical BIR2/IBM interface, Hid engages additional regions of DIAP1 including the RING domain and thereby further stabilizes the interaction. Regarding the Rpr interactions, despite the fact that Rpr1 and Rpr2 share the limited BIR1 surface in the quaternary complex (Figure 8D-G), their binding is overall as strong or stronger than in the DIAP1/Rpr1/Rpr2 trimer (Figure 7).

Specifically, while the DIAP1/Rpr1 interaction remains about the same ((3/10/166) in the trimer to (3/8/171) in the quaternary complex), the interaction of DIAP1/Rpr2 increases from (0/7/104) in the trimeric complex to (1/8/161) in the quaternary complex (Figure 7B, Figure 8B; Suppl. Table S1). These observations are consistent with a redistribution that maintains or enhances total binding strength. Finally, the Rpr1/Rpr2 interface itself is dramatically reinforced in the quaternary assembly, rising from (0/0/36) in the trimer to (1/6/357) in the quaternary complex (Figure 7B, Figure 8B; Suppl. Figure S7, Suppl. Table S1). This dimeric interaction is also much stronger than that observed in the isolated homodimer (2/0/48) (Figure 2A). Taken together, these comparisons show that the quaternary complex maximizes DIAP1 neutralization by combining an extensive BIR2/Hid engagement, a significant RING/Hid interaction, stable and cooperative dual Rpr binding at BIR1, and a strongly reinforced Rpr1/Rpr2 dimer (Figure 8, Suppl. Figure S7, Suppl. Table S1). This architecture suggests that the quaternary assembly is the most effective configuration for inactivating DIAP1, thereby promoting caspase activation and triggering apoptosis.

Assembly of the quaternary DIAP1/Hid/Rpr1/Rpr2 complex induces a profound reorganization of DIAP1 relative to its apo state. When BIR1 is used as the anchor, both BIR2 and the RING domain undergo large rigid-body displacements, with centroid shifts of ∼46 Å and ∼41 Å, respectively, and reorientation angles of ∼45-61° (Figure 8H). These repositionings suggest that in the fully engaged quaternary complex, the geometry of the RING domain is reconfigured in a way that could alter its functional specificity. Rather than optimally positioning itself for caspase ubiquitylation, the RING may be redirected toward promoting autoubiquitylation of DIAP1.

### 3.9 Structural and functional interaction of dBruce with Reaper

In addition to DIAP1, another IAP family member, dBruce, has been genetically identified as a potent inhibitor of Rpr-induced apoptosis (Arama *et al*., 2003; Vernooy *et al*., 2002). dBruce is a massive protein comprised of 4,876 aa, and contains a BIR domain at the N-terminus (aa 251-321) and a C-terminal Ubc domain (aa 4,636-4,790) with E2-ubiquitin-conjugating activity. The Ubc domain is thought to mediate non-canonical ubiquitination of Rpr (Domingues and Ryoo, 2012; Hao et al., 2004). We successfully modeled the full-length structure of dBruce, incorporating proper coordination of a Zn² ion in the BIR domain (Figure 9A). The model predicts a large central core composed of almost 200 high-confidence α-helices (pLDDT >90), connected by flexible, low-confidence loops. The PAE plot shows a mosaic of high- and low-confidence regions (Figure 9A), indicative of local flexibility, while the global prediction remains notably reliable given the unusually large size of this protein, and thus provides a solid foundation for subsequent interaction analysis.

**Figure 9.**
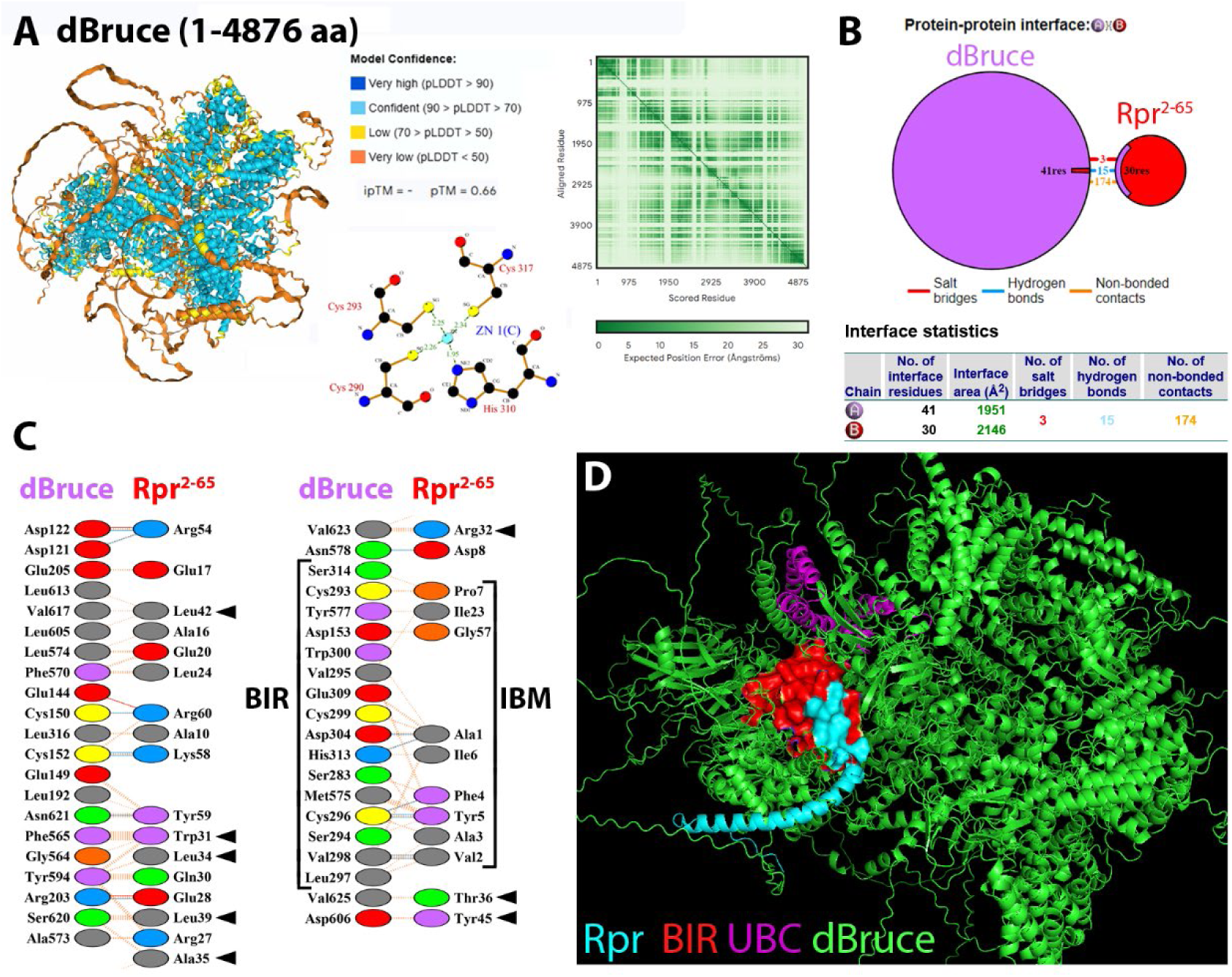
Structural model of full-length dBruce and its complex with Reaper^2-65^. (**A**) AF3 predicts that dBruce (4,876 aa) folds into a large core containing almost 200 high-confidence α-helices with unstructured loops connecting them. Coloring reflects pLDDT confidence according to the key. The PAE map indicates mixed global confidence throughout the protein. The BIR domain (residues 251-321) complexes one Zn^2+^ ion. (**B**) The dBruce/Rpr^2–65^ complex. Illustration of the protein-protein interface and interface statistics as obtained by PDBsum. (**C**) Specific residue interactions at the protein-protein interface. The type of interaction is colored coded according to the key. The IBM of Rpr binds the BIR domain of dBruce (brackets). Residues of the GH3 motif of Rpr^2-65^ (arrowheads) form a second interface outside the BIR domain of dBruce (involving residues between aa 564-625). (**D**) PyMOL-generated view of the dBruce/Rpr^2-65^ protein complex. Rpr^2-65^ is colored in cyan. dBruce is colored in green except for the BIR (red) and Ubc (magenta) domains. The IBM of Rpr^2-65^ and the BIR of dBruce are highlighted in surface mode.

Using AF3, we modeled the complex between dBruce and Rpr^2-65^ (Figure 9B,D).

Previous studies suggested that Rpr can bind the BIR domain of dBruce (Domingues and Ryoo, 2012; Vernooy *et al*., 2002). The GH3 domain of Rpr has also been implicated in the interaction with dBruce, but mechanistic details are unknown. The predicted dBruce/Rpr^2-65^ complex recapitulated key features of previously proposed interactions: the IBM of Rpr (Ala1, Ala3, Phe4, Tyr5, Ile6, Pro7, Asp8) formed contacts with several BIR residues in dBruce (Cys293, Val295, Trp300, Asp304, Glu309, His313, Ser314) (Figure 9C), consistent with prior biochemical data (Domingues and Ryoo, 2012). Sequence alignment of the BIR domain of dBruce with BIR1 and BIR2 of DIAP1 revealed conservation of key IBM-binding residues (Table 5), providing functional support for the prediction that dBruce engages the IBM of Rpr through a canonical BIR/IBM interaction. The dominant forces at the BIR/IBM interface are hydrophobic and backbone-directed hydrogen bonds, while electrostatic contributions play only a minor role.

**Table 5:**
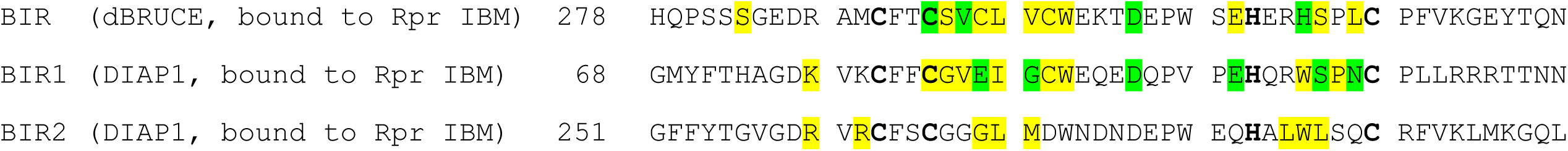
Comparisons of residues of BIR domains of dBruce and DIAP1 engaged with the IBM of Rpr. (see also Figure 9 and Supplementary Table S1) Sequence alignment reveals partial conservation between the BIR domain of dBRUCE (top) and the BIR1 and BIR2 domains of DIAP1. Green-highlighted residues of the aligned sequences form hydrogen bonds and non-bonded contacts with the IBM motif of Rpr, while yellow-highlighted residues indicate non-bonded contacts with this IBM. Despite sequence differences, the BIR domain interactions with the Rpr IBM display striking similarity. Bold residues (Cys, His) are chelating Zn^2+^ and are used as anchors for the alignment of the BIR domains.

Importantly, beyond this core interaction, the model also revealed a strong and extensive interface between the α-helical region of Rpr including the GH3 motif (Trp31, Leu34, Glu28, Arg32, Ala35, Tyr45) and a region of dBruce outside the BIR domain (residues 594-623) (Figure 9C). There is no direct engagement of Rpr with the UBC domain (Figure 9C,D). Therefore, the overall binding mode is consistent with a strong, specific BIR/IBM interaction, with auxiliary GH3 contacts likely enhancing complex stability or orienting the complex (Figure 9B,D).

These AF3-generated results confirm a previous model in which dBruce acts as a high-affinity sink for Rpr, sequestering its IBM motif and thereby blocking its ability to inactivate DIAP1 or engage Dronc (Domingues and Ryoo, 2012). This model explains well the genetic evidence identifying dBruce as an inhibitor of Rpr-induced apoptosis (Arama et al., 2007; Vernooy *et al*., 2002) and positions it as a potent endogenous regulator of apoptotic signaling.

## 4. Discussion

In this study, we present a comprehensive structural and functional characterization of the *Drosophila* IAP antagonists Reaper (Rpr), Hid, Grim, Sickle, and Jafrac2, with particular focus on their interactions among themselves and with DIAP1. Leveraging AlphaFold3 (AF3), we modeled the full-length conformations and multimeric complexes of these proteins, providing unprecedented insights into their functional interfaces and mechanistic roles in the regulation of apoptosis.

Our findings significantly expand current knowledge of the apoptotic pathway in *Drosophila* by resolving the full-length structures of the IAP antagonists and their target DIAP1 in both monomeric forms and multimeric complexes. Consistent with previous reports (Sandu *et al*., 2010), we find that Rpr is a structurally robust, dimerizing protein with a central α-helical domain that stabilizes its multimeric form and promotes its mitochondrial localization and apoptotic function. Hid, in contrast, lacks a well-resolved 3D structure. Aside from a small number of α-helices, the majority of Hid appears unstructured in our AF3 models. While this might initially seem like a limitation, we speculate that this structural plasticity is in fact advantageous, enabling Hid to flexibly adopt conformations required for binding diverse partners, including Rpr, Grim, Sickle and DIAP1.

Notably, the C-terminal tail-anchor (TA) helix of Hid plays a key role in recruiting Rpr to the outer mitochondrial membrane (OMM). This recruitment is essential for the formation of higher-order protein complexes and initiation of the apoptotic program. We speculate that specific interactions between Rpr and residues in the TA domain of Hid may serve as a targeting mechanism to direct Rpr to the OMM. Such physical proximity could promote the cooperative binding of IAP antagonists to DIAP1 and amplify apoptotic signaling.

A major finding of this study is the dual role of the N-terminal methionine (Met1) of Rpr.

While Met1 must be removed to expose the IBM of Rpr for DIAP1 binding (Wu *et al*., 2000), our models show that it actually stabilizes the Rpr/Hid complex by forming key interactions with residues 161-164 of Hid. Deletion of Met1 weakens the overall Rpr/Hid interaction, but exposes the IBM for apoptotic function. These findings reveal a previously unrecognized dual regulatory function of Met1 that establishes a novel checkpoint mechanism in the apoptotic pathway. While Met1 removal is essential for IBM/DIAP1 interaction and apoptosis, Met1 simultaneously serves a critical stabilizing function for the Rpr/Hid complex. This regulatory tradeoff creates an inherent temporal constraint where the modification (removal of Met1) enabling apoptosis also reduces stability of the complexes driving the apoptotic signal. This paradoxical relationship indicates that apoptotic activation is subject to self-limiting constraints that may prevent excessive cell death.

Equally important is our modeling of full-length DIAP1, which reveals a compact, globular architecture in which the BIR1, BIR2, and RING domains are tightly packed, rather than a modular, “beads-on-a-string” organization. Zn^2+^ ions play a critical role in maintaining this structure, coordinating with conserved cysteine and histidine residues to stabilize both BIR and RING domains.

Consistent with previous studies (Tenev *et al*., 2005; Wu *et al*., 2001; Zachariou *et al*., 2003), our models confirm that the IBMs of most IAP antagonists preferentially bind BIR2 of DIAP1. However, Rpr is an exception. While its IBM peptide alone also favors BIR2, the full-length Rpr^2-65^ protein binds strongly to BIR1. This binding specificity is mediated by α-helical residues in the Rpr backbone, most notably Glu20 which directly guides the IBM of Rpr toward BIR1, overriding the default BIR2 preference.

Nevertheless, a single residue in the IBM, Pro3, can reverse this specificity. Substitution of Ala3 with Pro in Rpr redirects binding from BIR1 to BIR2. Pro3 introduces steric hindrance with Trp103 of BIR1, but stabilizes contacts with Gly269, Leu270 and Trp286 in BIR2. These results underscore that BIR-binding specificity is not determined solely by residues in the backbone and the central α-helical regions, but also by cooperative effects from the IBM.

Our models of higher-order complexes also provide insights into a potential molecular mechanism by which IAP antagonists switch the E3-ubiquitin ligase activity of the RING domain of DIAP1. In the absence of IAP antagonists, the RING domain functions as a ubiquitin ligase that suppresses apoptosis by ubiquitylating caspases (Kamber Kaya et al., 2017; Wilson et al., 2002). Upon binding of IAP antagonists, particularly Rpr and Hid, the E3 ligase activity of the RING domain shifts toward auto-ubiquitylation and degradation of DIAP1 (Hays et al., 2002; Holley et al., 2002; Ryoo et al., 2002; Yoo et al., 2002), and the release of active caspases. The higher-order complexes discussed in this work provide a glimpse about how this switch in E3 ligase activity might happen.

Comparison of the structural effects of Rpr and Hid binding to DIAP1 reveals that these two IAP antagonists exploit distinct allosteric mechanisms to redirect the E3 ubiquitin ligase activity of the RING domain. Binding of Rpr to the BIR1 domain drives a large-scale reorientation of the RING domain toward BIR1, bringing the E3 ubiquitin ligase into proximity with the BIR1/IBM interface. This conformational change is well positioned to facilitate DIAP1 self-ubiquitylation and degradation, while simultaneously displacing bound caspases. In contrast, Hid binds via its IBM to BIR2, where the BIR2/RING module remains rigid, but BIR1 is displaced outward. Although this alignment does not reposition the RING toward BIR1, Hid makes direct contacts with RING residues through its central α-helix 2. These interactions suggest that Hid can still bias the RING domain toward autoubiquitylation, but by directly modulating RING contacts rather than by altering its domain orientation. Thus, both Reaper and Hid converge on the same functional endpoint, the self-destruction of DIAP1, but they achieve it through structurally divergent routes, underscoring the versatility of IAP antagonists in disabling DIAP1 function.

While the trimeric DIAP1/Rpr/Rpr complex demonstrates functional inhibition, the weaker individual binding interfaces and reduced Rpr dimer stability suggest this configuration is not optimal for sustained DIAP1 neutralization (Figure 7). In contrast, the quaternary DIAP1/Hid/Rpr1/Rpr2 complex represents the most effective architecture for DIAP1 inhibition through cooperative binding mechanisms. This assembly features dramatically enhanced interface strengths, including a reinforced DIAP1/Hid interaction that engages both BIR2 and RING domains, maintained or improved Rpr binding despite shared BIR1 occupancy, and a substantially strengthened Rpr1/Rpr2 interface facilitated by direct IBM-IBM contacts (Figure 8; Suppl. Figure S7). Importantly, quaternary complex formation induces profound DIAP1 reorganization, with the RING domain undergoing a ∼41 Å centroid shift and ∼45-61° reorientation relative to the apo state (Figure 8H). We speculate that such rearrangements could switch the activity of the RING E3 ligase from caspase ubiquitination to auto-ubiquitylation of DIAP1, thereby promoting its degradation and amplifying the apoptotic signal. Consistently, a reorganization of the RING domain upon antagonist binding has also been observed for cIAP2 (Dueber et al., 2011).

Jafrac2, a thioredoxin peroxidase localized to the ER under normal conditions, represents a structurally distinct class of IAP antagonists (Tenev *et al*., 2002). Unlike the other IAP antagonists that act at mitochondria and serve as dedicated apoptosis regulators, Jafrac2 performs essential oxidative folding functions in the ER. Upon apoptotic signaling or stress, mature Jafrac2^18–242^ is released into the cytosol, where its IBM binds preferentially to BIR2 of DIAP1.

Our modeling shows that the peroxiredoxin domain stabilizes this interaction by contacting BIR1 and the RING domain. These findings demonstrate how proteins with non-apoptotic roles can acquire death-promoting functions through proteolytic processing and re-localization.

Finally, we modeled the full-length structure of dBruce, a massive IAP family member with almost 200 α-helices and a compact core. dBruce strongly interacts with Rpr via a canonical BIR/IBM interface, and forms additional contacts involving the GH3 motif of Rpr (Figure 9).

This inhibitory complex not only sequesters Rpr from DIAP1, but may also redirect Rpr for degradation. Remarkably, dBruce has been shown to ubiquitinate Rpr via an unconventional mechanism, possibly targeting the N-terminal amino group rather than lysine residues (Domingues and Ryoo, 2012). This highlights a unique regulatory mechanism whereby an IAP family member suppresses apoptosis not by blocking caspases, but by selectively degrading IAP antagonists. This mode of apoptotic regulation is analogous to that employed by viral IAP proteins (Vucic *et al*., 1997; Vucic *et al*., 1998).

Together, these structural insights significantly advance our understanding of the molecular mechanisms underlying apoptosis in *Drosophila*. They reveal how molecular determinants such as Met1, Pro3, Glu20, GH3 motifs, zinc coordination and flexible regions orchestrate precise interactions between IAPs and their antagonists. They also illustrate how oligomerization and dynamic multimeric complexes tune the strength, timing, and outcome of the apoptotic response. Given the evolutionary conservation of these pathways, the principles uncovered here likely also have broader implications for understanding apoptosis in mammals, where homologous proteins such as Smac, XIAP, and BRUCE/Apollon play similar roles.

We acknowledge several limitations of this work. All conclusions are derived from computational models and lack experimental validation through high-resolution structural methods such as cryo-EM or X-ray crystallography. However, all previously established interactions, including the BIR domain preferences of Rpr, Hid, and Grim, and the Zn^2+^-binding architecture of DIAP1, were accurately recapitulated by AF3. This concordance lends confidence to our novel predictions, particularly those involving full-length proteins in higher order complexes.

In conclusion, our study demonstrates the remarkable potential of AlphaFold3 to elucidate intricate protein-protein interactions and uncover fundamental principles of apoptotic regulation in *Drosophila*. By integrating structural predictions with mechanistic analysis, we reveal how spatial and temporal control of cell death emerges from a dynamic network of competing molecular interactions, adding new dimensions to our understanding of apoptosis regulation. These findings not only refine current models of caspase control but also open new avenues for exploring non-apoptotic and evolutionarily conserved roles of apoptotic proteins. Looking ahead, the predictive power of deep-learning-based structure modeling promises to accelerate the discovery of novel regulatory interactions across diverse signaling pathways, ultimately bridging structural insights with physiological function in development, homeostasis, and disease.

## Acknowledgments

This work was supported by a grant from the National Institute of General Medical Sciences (NIGMS) under award number R35GM118330 to AB. The content is solely the responsibility of the authors and does not necessarily represent the official views of the NIH.

## Author contributions

PR: Conceptualization, Data curation, Formal Analysis, Investigation, Methodology, Validation, Writing–review and editing.

AB: Conceptualization, Data curation, Formal Analysis, Funding acquisition, Resources, Supervision, Writing–original draft, Writing–review and editing.

---

Supplementary Figure S1. Dimer models of Hid, Grim, Sickle, and Jafrac2. (related to Figure 2)

AF3-predicted models of the dimers of Hid (A), Grim (B), Sickle (C) and Jafrac2 (D). Confidence values vary across interfaces and are low except for the Jafrac2 dimer.

Supplementary Figure S2. Heterodimeric interactions between IAP antagonists. (related to Figure 2)

AF3-predicted models of (A) Hid/Grim, (B) Rpr/Grim, and (C) Hid/Sickle heterodimeric complexes. The a1, a2, b1, b2, c1 and c2 panels illustrate the protein-protein interfaces, interface statistics and residue interactions at the interfaces of the heterodimers according to PDBsum. The a3-a5, b3-b5 and c3-c5 panels show PyMOL-generated structures of the heterdimeric complexes. Hid and Grim (b4) are colored in green with the interacting residues highlighted in yellow. Rpr, Grim (a4) and Sickle are shown in cyan with the interacting residues highlighted in red.

Supplementary Figure S3. IBM peptide interactions with isolated BIR2. (related to Figure 4) (A-D) PDBsum- and PyMOL-generated images of the interaction of the IBM peptides (10mers) of Rpr (A), Hid (B), Grim (C) and Sickle (D) with the isolated BIR2 domain (residues 201 to 325) of DIAP1.

(a1, b1, c1, d1) The PDBsum-generated diagrams present a 2D interaction map with the IBM peptide shown in the center. IBM residues are labeled in blue, while residues from the BIR2 domain that engage in hydrogen bonding with the peptide are labeled in green. Hydrogen bonds are indicated as green dashed lines. Hydrophobic contacts are depicted as spoked arcs (half-circles with radiating lines), drawn around BIR2 residues whose side chains form non-bonded interactions with IBM residues. A complete list of all hydrogen bonds and non-bonded contacts is provided in Supplementary Table S1.

(a2, a3, b2, b3, c2, c3, d2, d3) PyMOL-rendered images depicting the interaction between the IBM peptides and the BIR2 domain. The BIR2 domain is shown in yellow, the IBM peptide in cyan. Hydrogen bonds are in magenta, while non-bonded contacts are not displayed. The lower panels indicate the specific BIR2 residues forming hydrogen bonds with IBM residues. These interactions closely mirror those reported in previously characterized IBM/BIR2 complexes of Reaper, Hid, and Grim (Wu *et al*., 2001) and are summarized in Table 2.

Supplementary Figure S4. IBM peptides of all IAP antagonists interact preferentially with the BIR2 of full-length DIAP1. (related to Figure 4)

(A-D) PDBsum- and PyMOL-generated images of the interaction of the IBM peptides (10mers) of Rpr (A), Hid (B), Grim (C) and Sickle (D) with full-length DIAP1. All IBM peptides preferentially bind BIR2 in full-length DIAP1 context. Shared BIR2 contact residues are Gly269, Met271, Asp277, Gln282, Trp286.

(a1, b1, c1, d1) The PDBsum-generated diagrams present a 2D interaction map with the IBM peptide shown in the center. IBM residues are labeled in blue, while residues from the BIR2 domain that engage in hydrogen bonding with the peptide are labeled in green. Hydrogen bonds are indicated as green dashed lines. Hydrophobic contacts are depicted as spoked arcs (half-circles with radiating lines), drawn around the BIR2 residues whose side chains form non-

bonded interactions with IBM residues. A complete list of all hydrogen bonds and non-bonded contacts is provided in Supplementary Table S1. (a2, a3, b2, b3, c2, c3, d2, d3) PyMOL-rendered images depicting the interaction between the IBM peptides and the BIR2 domain in full-length DIAP1. The domains are colored according to the following code: BIR1 - red; BIR2 - yellow; RING - magenta; IBM peptide - cyan. The lower panels indicate the specific BIR2 residues forming hydrogen bonds (magenta) with IBM residues. These interactions closely mirror those reported in previously characterized IBM/BIR2 complexes of Reaper, Hid, and Grim (Wu *et al*., 2001) and are summarized in Table 3.

Supplementary Figure S5. Interaction of full-length DIAP1 with Grim^2-138^ and Sickle^2-108^.

(related to Figure 4)

(A) The DIAP1/Grim^2-138^ complex. (a1) Illustration of the protein-protein interface and interface statistics as obtained by PDBsum. (a2) Specific residue interactions at the protein-protein interface. The IBM of Grim^2–138^ binds to BIR2 of DIAP1 (see brackets). (b3) Overall view of the DIAP1/Grim^2–138^ complex. The individual domains are colored as indicated. Linker regions are omitted for clarity. The white arrow points to the BIR2/IBM interface. (b4) Detailed interaction view of the BIR2/IBM interaction. Critical residues for this interaction are labeled.

(B) The DIAP1/Sickle^2-108^ complex. (b1) Illustration of the protein-protein interface and interface statistics as obtained by PDBsum. (b2) Specific residue interactions at the protein-protein interface. The IBM of Sickle^2–108^ binds to BIR2 of DIAP1 (see brackets). Sickle^2–108^ also uses residues of its α-helix (aa 70-84) and further C-terminal residues to contact the RING domain of DIAP1 (arrowheads). (b3) Overall view of the DIAP1/Sickle^2-108^ complex. The individual domains are colored as indicated. Linker regions are omitted for clarity. The white arrow points to the BIR2/IBM interface. (b4) Detailed interaction view of the IBM/BIR2 interaction. Critical residues for this interaction are labeled.

Supplementary Figure S6. PyMOL-generated images of the protein complexes consisting of DIAP1 and IAP antagonist chimera or Rpr point mutants. (related to Figure 5)

In each case, the modeling was done with full-length DIAP1. The IAP antagonist chimera and Rpr point mutants are indicated above each panel. The coloring is as follows: DIAP1 - BIR1 in red, BIR2 in yellow, RING in magenta; IAP antagonists are in blue except for the IBM which is cyan. The interactions interfaces are shown in surface mode. Zn^2+^ ions are orange. White arrows in each panel point to the BIR/IBM interface.

Supplementary Figure S7. Missing panels of Figure 8. (related to Figure 8)

(A) Illustration of the protein-protein interfaces and interface statistics as obtained by PDBsum.

(B,C) Specific residue interactions at the interface of Hid and Rpr2 (B), and of Rpr1 and Rpr2

(C) as obtained by PDBsum. The two IBM motifs of Rpr1 and Rpr2 interact with each other.

Supplementary Table S1. Complete list of all atom-atom interactions across the protein/protein interfaces in the protein complexes described in this paper, as revealed by PDBsum. (related to Figures 2, 4, 5, 6, 7, 8, 9, S2, S3, S4, S5) Listed are the specific atom-atom interactions which form hydrogen bonds, non-bonded contacts and salt bridges. The cutoff for non-bonded contacts is set as <4.0 Angstroms.

