## Supplemental Figure S1-S5 for "AlphaFold3-based modeling uncovers the dynamic structural interface between full-length IAP antagonists and DIAP1 for apoptosis regulation in *Drosophila*"

Suppl Fig S1

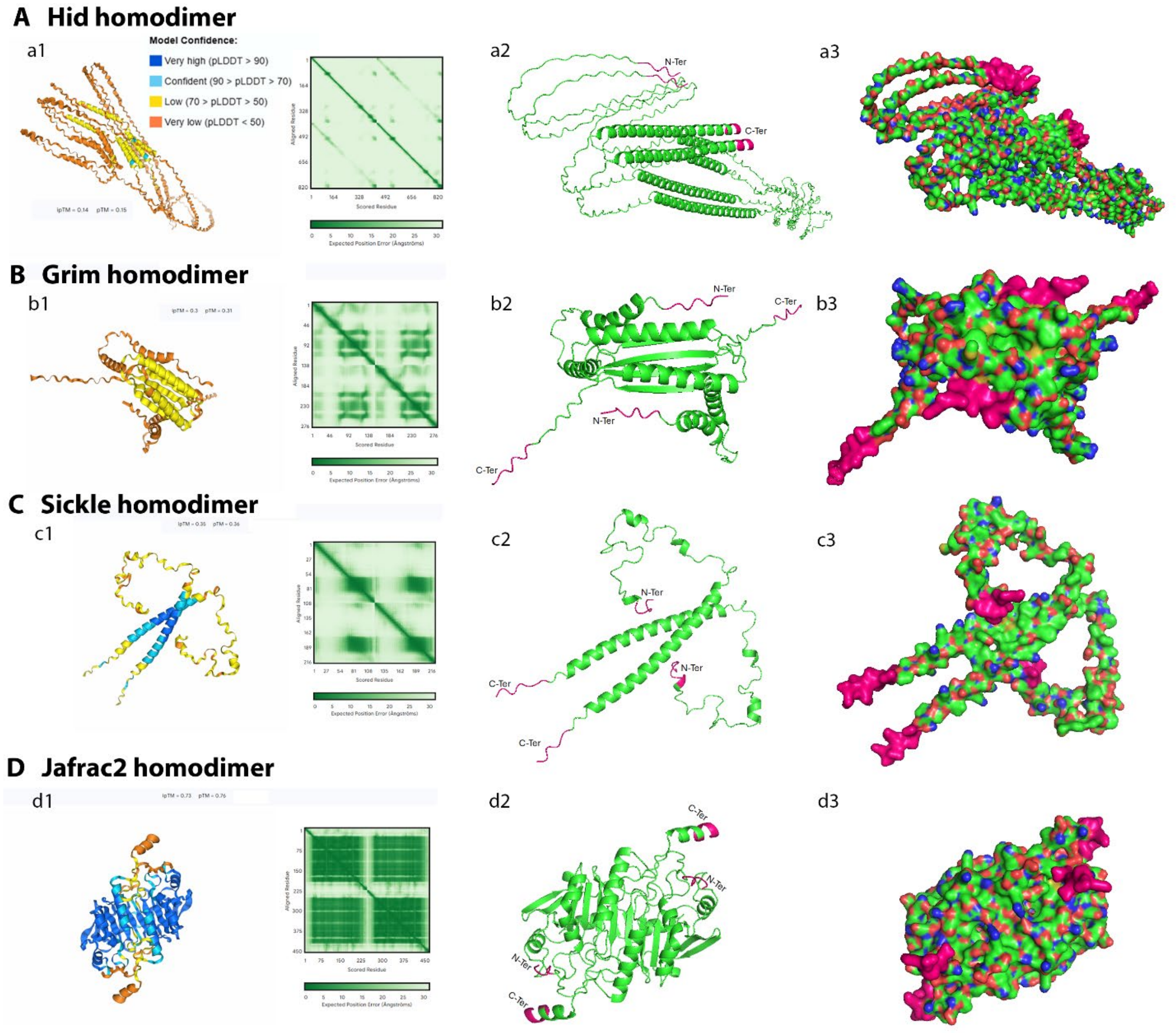
A Hid homodimer
a1
Model Confidence:
Very high (pLDDT > 90)
Confident (90 > pLDDT > 70)
Low (70 > pLDDT > 50)
Very low (pLDDT < 50)
ipTM = 0.14 pTM = 0.15
Aligned Residue
Scored Residue
Expected Position Error (Ångströms)
a2
N-Ter
C-Ter
a3
B Grim homodimer
b1
ipTM = 0.3 pTM = 0.31
b2
N-Ter
C-Ter
N-Ter
C-Ter
b3
C Sickle homodimer
c1
ipTM = 0.35 pTM = 0.36
c2
N-Ter
N-Ter
C-Ter
C-Ter
c3
D Jafrac2 homodimer
d1
ipTM = 0.73 pTM = 0.76
d2
C-Ter
N-Ter
N-Ter
C-Ter
d3

Suppl Fig S2

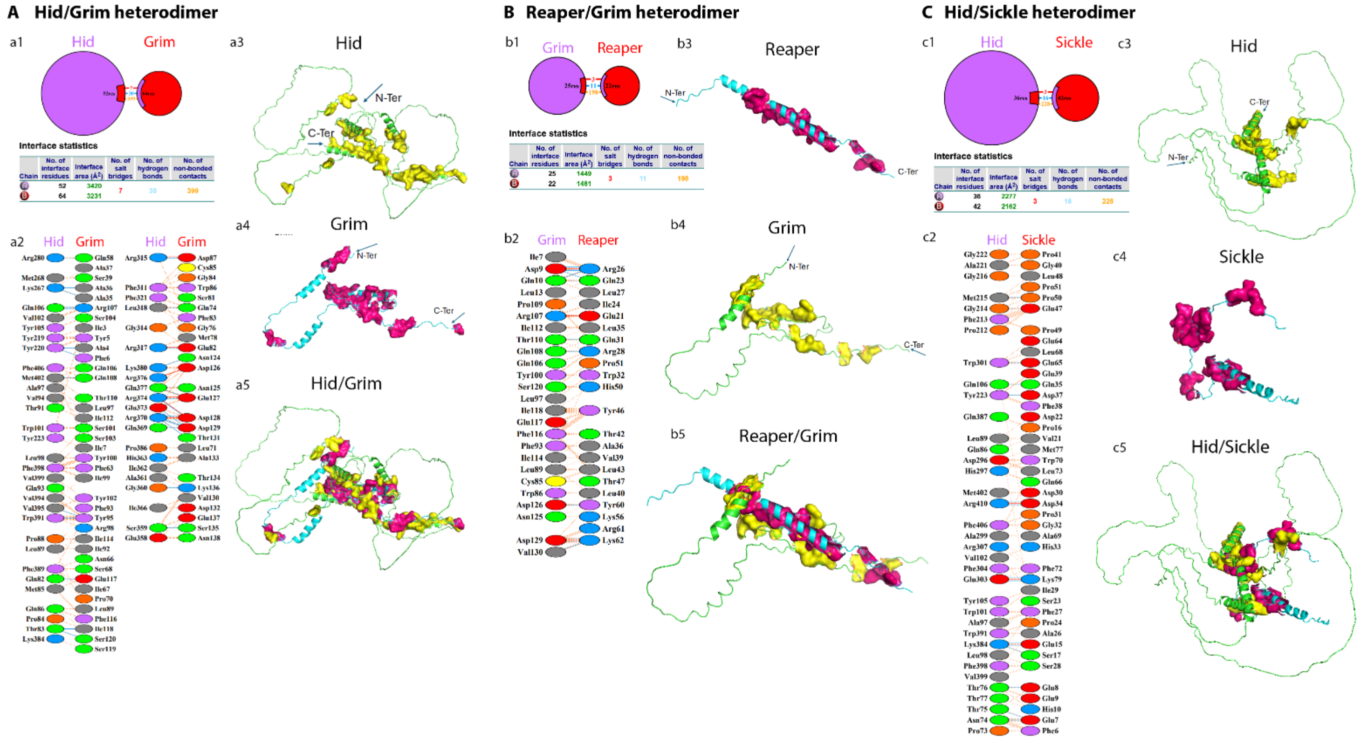
A Hid/Grim heterodimer
a1
Hid
Grim
Interface statistics
a2
Hid
Grim
Hid
Grim
a3
Hid
N-Ter
C-Ter
a4
Grim
N-Ter
C-Ter
a5
Hid/Grim
B Reaper/Grim heterodimer
b1
Grim
Reaper
Interface statistics
b2
Grim
Reaper
b3
Reaper
N-Ter
C-Ter
b4
Grim
N-Ter
C-Ter
b5
Reaper/Grim
C Hid/Sickle heterodimer
c1
Hid
Sickle
Interface statistics
c2
Hid
Sickle
c3
Hid
C-Ter
N-Ter
c4
Sickle
c5
Hid/Sickle

Suppl Fig S3

**A** Reaper IBM Peptide (2-11) in complex with BIR2

Ala-Val-Ala-Phe-Tyr-Ile-Pro-Asp-Gln-Ala

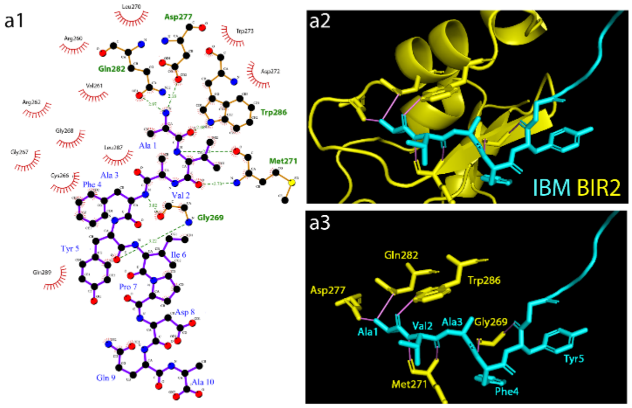


**B** Hid IBM Peptide (2-11) in complex with BIR2

Ala-Val-Pro-Phe-Tyr-Leu-Pro-Glu-Gly-Gly

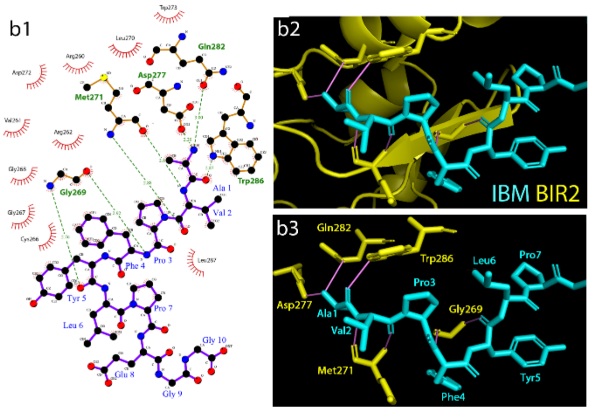


**C** Grim IBM Peptide (2-11) in complex with BIR2

Ala-Ile-Ala-Tyr-Phe-Ile-Pro-Asp-Gln-Ala

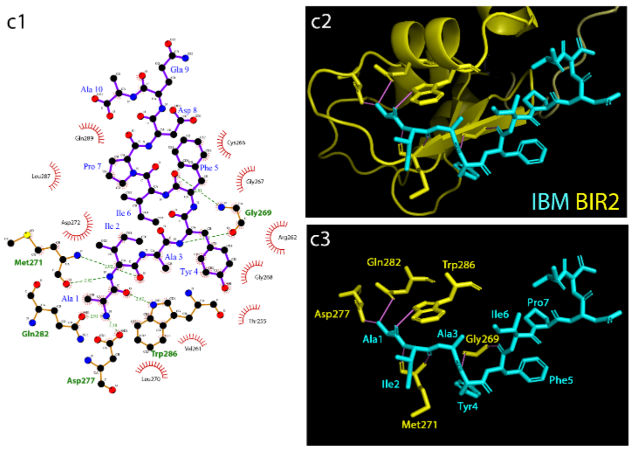


**D** Sickle IBM Peptide (2-11) in complex with BIR2

Ala-Ile-Pro-Phe-Phe-Glu-Glu-Glu-His-Ala

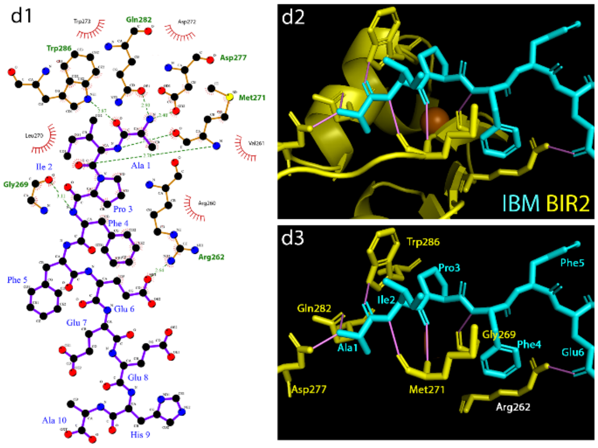

Suppl Fig S4

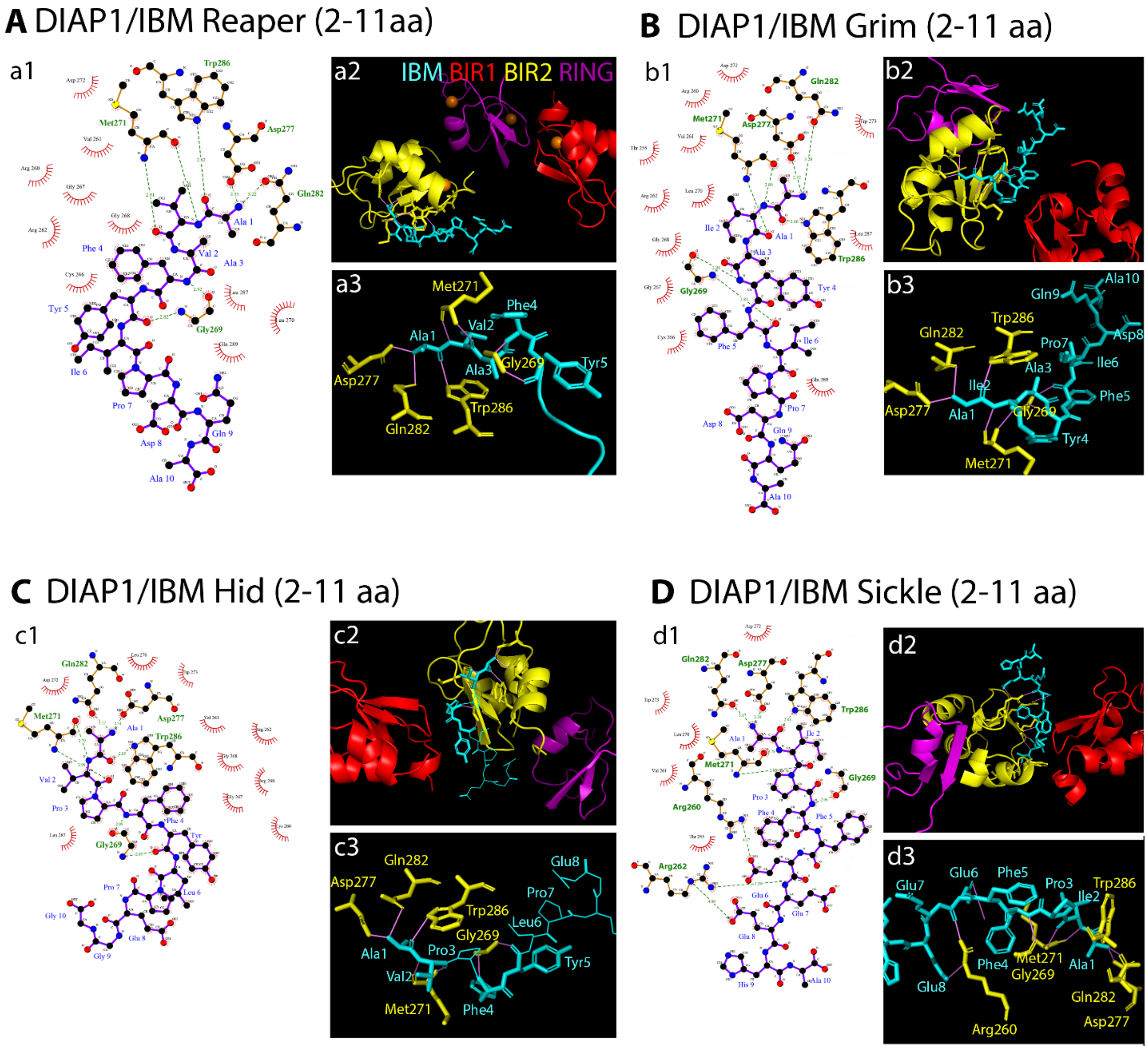
A DIAP1/IBM Reaper (2-11aa)
a1
a2
IBM BIR1 BIR2 RING
a3
Met271
Phe4
Val2
Ala1
Asp277
Ala3
Gly269
Tyr5
Trp286
Gln282
B DIAP1/IBM Grim (2-11 aa)
b1
b2
b3
Gln9
Ala10
Trp286
Gln282
Pro7
Asp8
Ile6
Ala3
Ile2
Asp277
Ala1
Gly269
Phe5
Tyr4
Met271
C DIAP1/IBM Hid (2-11 aa)
c1
c2
c3
Gln282
Glu8
Asp277
Pro7
Trp286
Leu6
Gly269
Ala1
Pro3
Tyr5
Val2
Met271
Phe4
D DIAP1/IBM Sickle (2-11 aa)
d1
d2
d3
Glu7
Glu6
Phe5
Pro3
Trp286
Ile2
Phe4
Met271
Gly269
Ala1
Glu8
Gln282
Arg260
Asp277

Suppl Fig S5

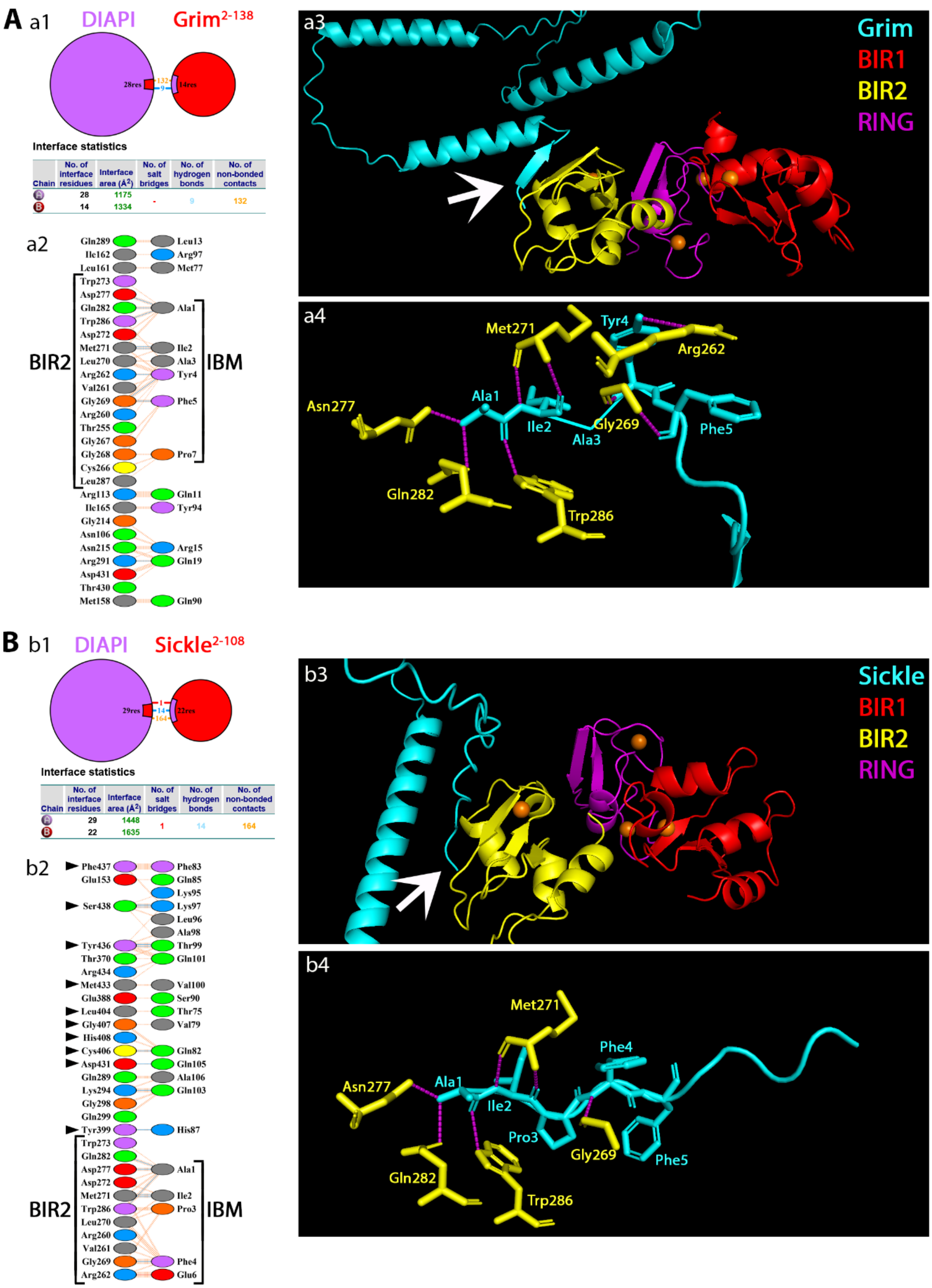
A
a1
DIAPI
Grim2-138
28res
132
9
14res
Interface statistics
Chain | No. of interface residues | Interface area (Å2) | No. of salt bridges | No. of hydrogen bonds | No. of non-bonded contacts
A | 28 | 1175 | - | 9 | 132
B | 14 | 1334
a2
Gln289 Leu13
Ile162 Arg97
Leu161 Met77
Trp273
Asp277
Gln282 Ala1
Trp286
Asp272
Met271 Ile2
Leu270 Ala3
Arg262 Tyr4
Val261
Gly269 Phe5
Arg260
Thr255
Gly267
Gly268 Pro7
Cys266
Leu287
Arg113 Gln11
Ile165 Tyr94
Gly214
Asn106
Asn215 Arg15
Arg291 Gln19
Asp431
Thr430
Met158 Gln90
BIR2
IBM
a3
Grim
BIR1
BIR2
RING
a4
Met271
Tyr4
Arg262
Asn277
Ala1
Ile2
Gly269
Ala3
Phe5
Gln282
Trp286
B
b1
DIAPI
Sickle2-108
29res
1
14
164
22res
Interface statistics
Chain | No. of interface residues | Interface area (Å2) | No. of salt bridges | No. of hydrogen bonds | No. of non-bonded contacts
A | 29 | 1448 | 1 | 14 | 164
B | 22 | 1635
b2
Phe437 Phe83
Glu153 Gln85
Lys95
Ser438 Lys97
Leu96
Ala98
Tyr436 Thr99
Thr370 Gln101
Arg434
Met433 Val100
Glu388 Ser90
Leu404 Thr75
Gly407 Val79
His408
Cys406 Gln82
Asp431 Gln105
Gln289 Ala106
Lys294 Gln103
Gly298
Gln299
Tyr399 His87
Trp273
Gln282
Asp277 Ala1
Asp272
Met271 Ile2
Trp286 Pro3
Leu270
Arg260
Val261
Gly269 Phe4
Arg262 Glu6
BIR2
IBM
b3
Sickle
BIR1
BIR2
RING
b4
Met271
Phe4
Asn277
Ala1
Ile2
Pro3
Gly269
Phe5
Gln282
Trp286

Suppl Fig S6

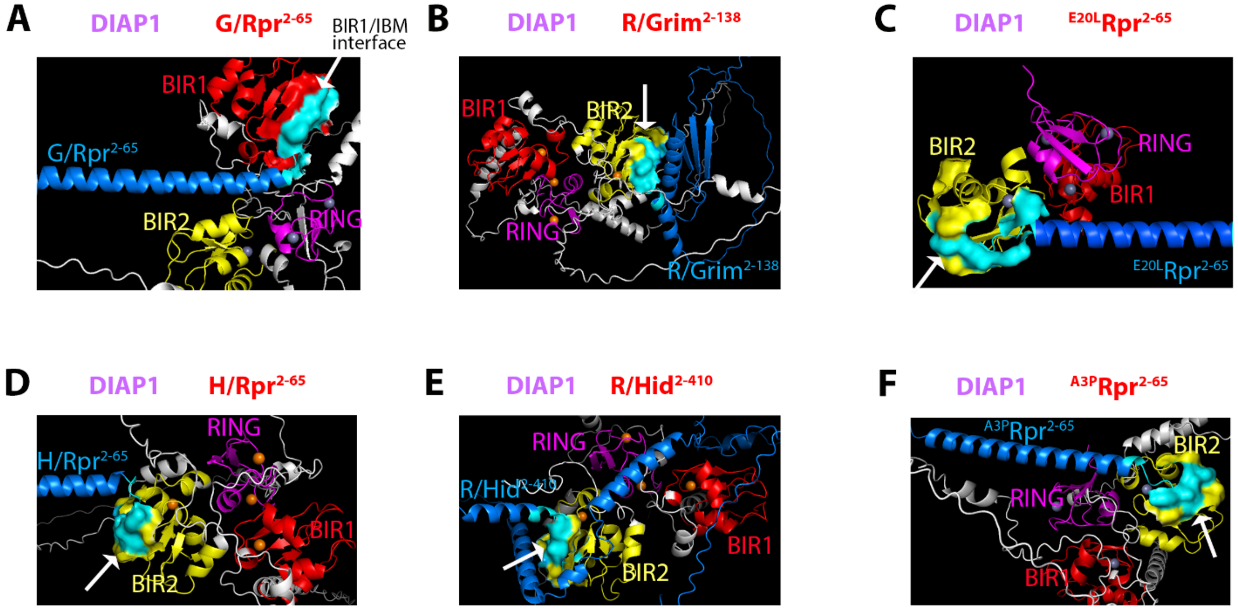

A
DIAP1
G/Rpr2-65
BIR1/IBM interface
BIR1
G/Rpr2-65
BIR2
RING
B
DIAP1
R/Grim2-138
BIR1
BIR2
RING
R/Grim2-138
C
DIAP1
E20LRpr2-65
BIR2
RING
BIR1
E20LRpr2-65
D
DIAP1
H/Rpr2-65
RING
H/Rpr2-65
BIR1
BIR2
E
DIAP1
R/Hid2-410
RING
R/Hid2-410
BIR1
BIR2
F
DIAP1
A3PRpr2-65
A3PRpr2-65
BIR2
RING

Suppl Fig S7

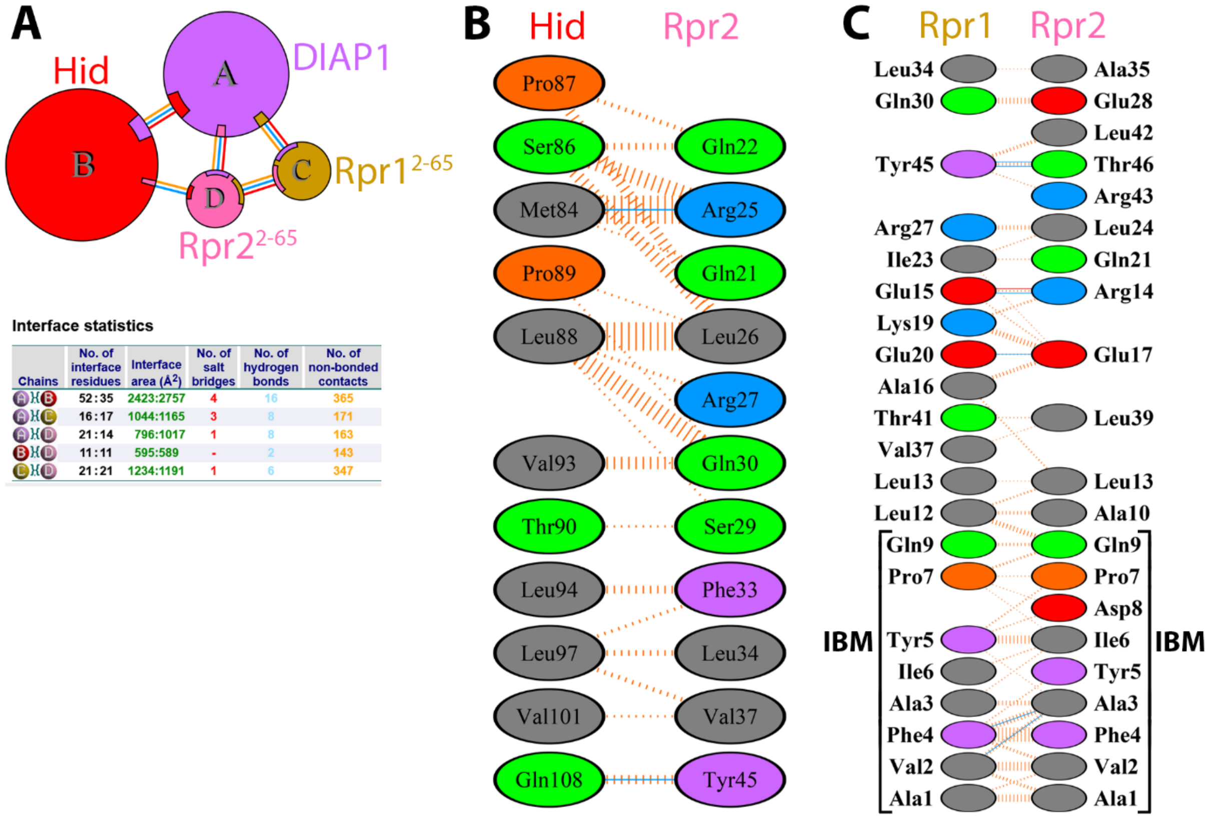


**Interface statistics**

| Chains | No. of interface residues | Interface area (Å$^2$) | No. of salt bridges | No. of hydrogen bonds | No. of non-bonded contacts |
|---|---|---|---|---|---|
| A–B | 52 : 35 | 2423:2757 | 4 | 16 | 365 |
| A–C | 16 : 17 | 1044:1165 | 3 | 8 | 171 |
| A–D | 21 : 14 | 796:1017 | 1 | 8 | 163 |
| B–D | 11 : 11 | 595:589 | - | 2 | 143 |
| C–D | 21 : 21 | 1234:1191 | 1 | 6 | 347 |
